# High-resolution structure of a microtubule-like tube composed of FtsZ–monobody complexes

**DOI:** 10.1101/2022.10.05.510932

**Authors:** Junso Fujita, Hiroshi Amesaka, Takuya Yoshizawa, Natsuko Kuroda, Natsuki Kamimura, Mizuho Hara, Tsuyoshi Inoue, Keiichi Namba, Shun-ichi Tanaka, Hiroyoshi Matsumura

## Abstract

FtsZ, a bacterial tubulin homologue, forms protofilaments and the Z-ring, which acts as a scaffold for accessory proteins during cell division. Although various studies have revealed its molecular mechanisms, the lack of high-resolution solution structures has hindered the understanding of the detailed mechanisms. Here, we developed a monobody (Mb) that binds FtsZs from *Escherichia coli* and *Klebsiella pneumoniae* (KpFtsZ) without affecting their GTPase activities. When expressed in *E. coli* cells, the Mb did not inhibit Z-ring formation but did inhibit cell division. The crystal structures of the KpFtsZ–Mb complexes revealed the epitope, and the cryoEM structure at 2.67 Å resolution showed a double helical tube consisting of two KpFtsZ protofilaments stabilized by the Mb filling interfilament gaps. Our structural analyses highlight the similarity between the microtubule and the FtsZ tube and the importance of the plasticity of FtsZ protofilaments.

## Introduction

FtsZ is a tubulin homologue GTPase protein that is widely conserved in bacteria and plays a key role in a complex called divisome that functions during cell division^1–3^. FtsZ polymerizes into protofilaments and a ring-shaped structure (Z-ring) with GTP, which is located in the middle of the cell and constricts to provide force for dividing the cell in half^4,5^. FtsZ is tethered to the cell membrane through other proteins such as FtsA and ZipA^6–8^, both of which recognize a flexible linker and C-terminal peptide of FtsZ and stabilize highly curved FtsZ protofilaments^9^. The mechanism of force generation has been discussed, and cell wall remodeling has become a primary candidate to produce this force, which is regulated by FtsZ treadmilling associated with its GTPase activity^10–14^.

To reveal the molecular mechanism leading to the FtsZ function, various crystal structures of FtsZ from different species have been determined^15–18^. Most of them show similar monomer conformations (relaxed, R conformation) and do not form protofilament-like structures in the crystals. FtsZ from *Staphylococcus aureus* (SaFtsZ) is the first species to exhibit a polymerization-preferred conformation (tense, T conformation) or both T and R conformations in filamentous structures in the crystals^19–22^. Recent studies have revealed filamentous crystal structures of FtsZ in the R conformation in two closely related species, *Escherichia coli* (EcFtsZ) and *Klebsiella pneumoniae* (KpFtsZ)^23,24^. As these crystal structures cannot escape the effect of crystal packing, trials have been conducted to determine the structure of FtsZ filaments in solution. Szwedziak *et al*. used cryo-electron tomography to observe reconstituted Z-rings in constricting liposomes^25^, and Wagstaff *et al*. reconstructed a low-resolution map of EcFtsZ protofilaments adopting the T conformation using cryo-electron microscopy (cryoEM) single particle analysis^26^. In addition, FtsZ forms minirings and thick helical tubes in the presence of specific support layers or accessary proteins^9,27-31^. However, due to the lack of high-resolution structures of the FtsZ protofilament in solution, the details of intramolecular and intermolecular interactions remain to be elucidated.

In this study, we developed a monobody (Mb), a synthetic binding protein based on the tenth human fibronectin type III domain (FN3)^32^ that binds to KpFtsZ and EcFtsZ to aid high-resolution structure determination. In *E. coli* cells, EcFtsZ complexed with this Mb could form Z-rings, but inhibits proper cell division to elongate the cells. We also determined the crystal structures of the R-conformation KpFtsZ–Mb and EcFtsZ–Mb complexes at a maximum resolution of 2.6–1.8 Å in two different space groups, in which the two protofilaments exhibited different curvatures. The negative staining EM observations showed that KpFtsZ formed various types of filaments depending on the additives. KpFtsZ alone can form thick and flexible tubes with a diameter of ∼25 nm, which is very similar to the helical tubes of other FtsZs^27–30^. These tubes were stabilized by the addition of the Mb. The cryoEM structure of the R-conformation KpFtsZ–Mb thick tube was determined at a resolution of 2.67 Å and showed a microtubule-like double helical tube composed of two KpFtsZ protofilaments stabilized by the Mb. The curvature of the protofilaments was much larger than that of the crystal structures. Our structural analyses showed the structural similarity and difference between the microtubule and FtsZ tube and the plasticity of the FtsZ protofilament, which may be important for the formation of Z-rings in various sizes for completing cell division.

## Results

### Selection and characterization of a monobody targeted to KpFtsZ and EcFtsZ

To generate monobodies that bind to KpFtsZ and EcFtsZ, we selected EcFtsZ using two combinatorial phage display libraries: the loop library and the side library (Fig. 1a). We used EcFtsZ as a target for this selection as EcFtsZ and KpFtsZ are highly homologous with a sequence identity of 98.7% (Fig. 1b). After four rounds of selection against EcFtsZ, five clones exhibiting specific binding to both KpFtsZ and EcFtsZ, as confirmed by ELISA (Extended Data Fig. 1), were selected for sequencing, yielding one unique clone Mb(Ec/KpFtsZ_S1) derived from the side library (Fig. 1c; for brevity, hereafter, we will use an abbreviated name for the monobody where the “Ec/KpFtsZ_” segment is omitted). Measurements of the dissociation constant (*K*_d_) in the yeast surface display format revealed that the Mb had *K*_d_ values in the single μM range to KpFtsZ and EcFtsZ (*K*_d_ = 5.40 ± 0.04 μM for KpFtsZ and *K*_d_ = 1.21 ± 0.34 μM for EcFtsZ; Fig. 1d). We then tested whether the addition of different types of nucleotides affected the binding of Mb(S1). The results showed that the binding did not change significantly in response to any of the nucleotides/nucleotide-analogues tested (Fig. 1e), and the binding of Mb(S1) to truncated KpFtsZ (residues 11–316; KpFtsZtr) was also confirmed by gel filtration chromatography (Extended Data Fig. 2), suggesting that the Mb binds to the GTPase domain of FtsZ.

**Fig. 1.**
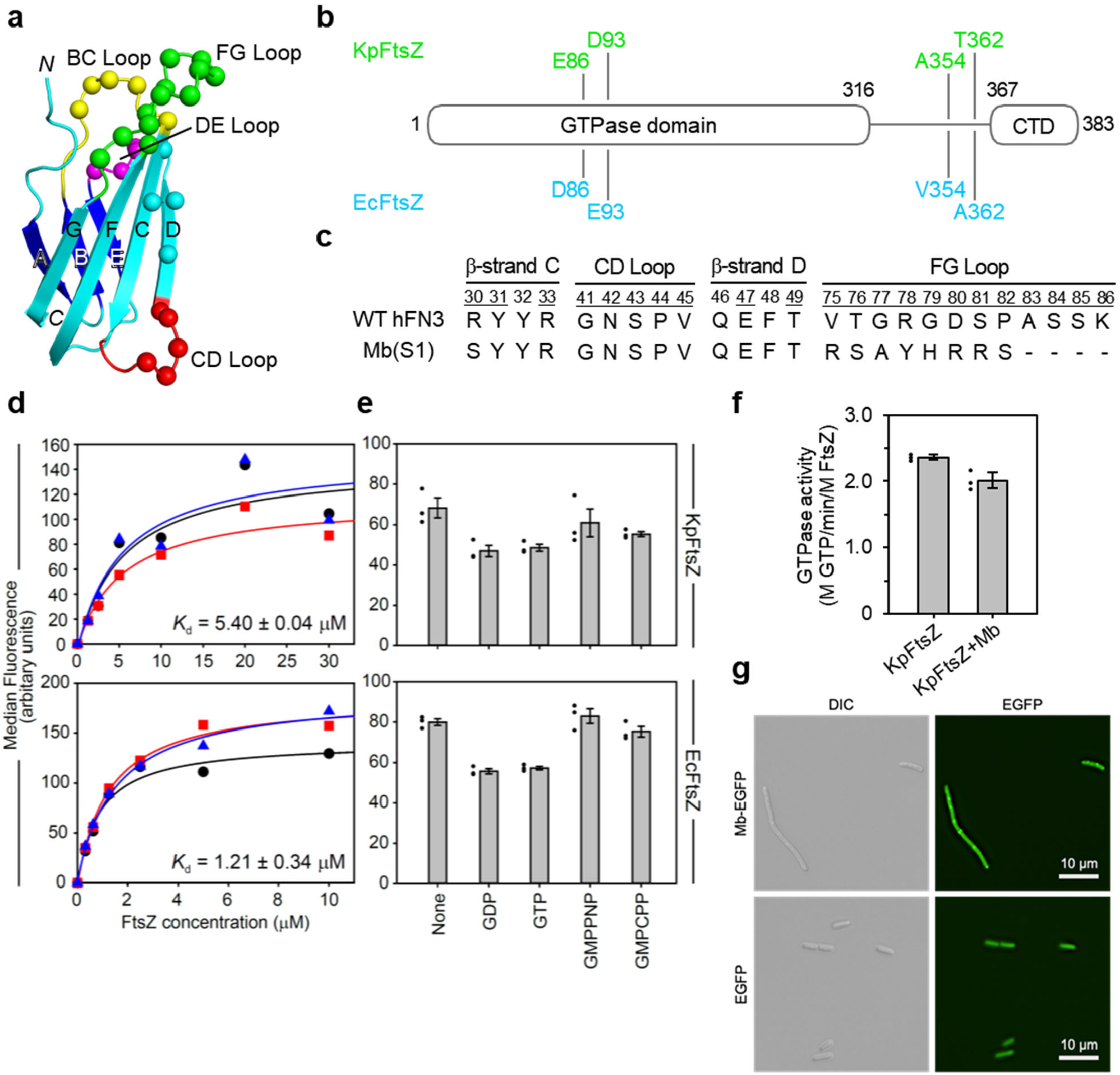
Monobody binding to KpFtsZ and EcFtsZ. **a**, Schematic of the monobody scaffold, with the locations of diversified residues shown as spheres and strands, loops and termini labeled. **b**, High sequence homology between EcFtsZ and KpFtsZ. Positions with different amino acid residues between the two FtsZs are indicated. **c**, Amino acid sequence of Mb(S1), with the wild-type hFN3 sequence as a reference. Residue numbers for diversified positions are underlined. **d**, Binding titration curves and the dissociation constants (*K*_d_) of Mb(S1) against KpFtsZ (upper panel) and EcFtsZ (lower panel) measured using yeast surface display. The median fluorescence intensities of yeast cells displaying Mb(S1) are plotted as a function of the concentration of KpFtsZ or EcFtsZ. The *K*_d_ values and errors shown are mean and standard deviations of three independent measurements. **e**, Effect of nucleotide/nucleotide-analogues on Mb(S1) binding to KpFtsZ (upper panel) or EcFtsZ (lower panel). **f**, GTPase activity assay of KpFtsZ with or without Mb(S1). Values are shown as the mean of three independent experiments with standard deviation. **g**, Differential interference contrast (DIC) and fluorescence microscopy images of *E. coli* cells overproducing EGFP and Mb(S1)-EGFP.

**Fig. 2.**
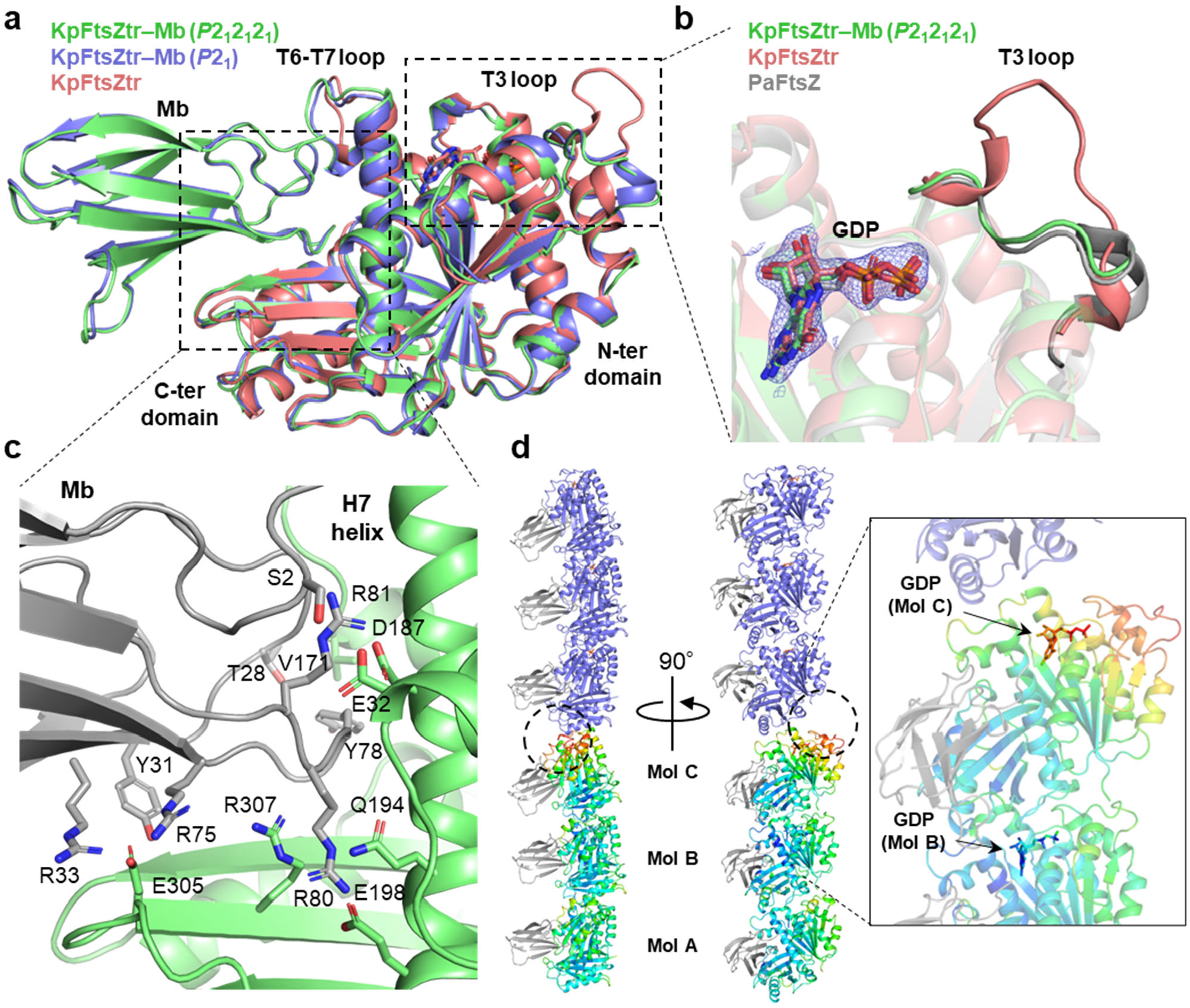
Crystal structures of KpFtsZtr–Mb complex. **a**, Superimposed crystal structures of the KpFtsZtr–Mb complex in the *P*2_1_2_1_2_1_ and *P*2_1_ space group and Mb-free KpFtsZtr (PDB code:6LL5). Only FtsZ moiety is used for superimposition. **b**, Close-up view around the T3 loop. Crystal structures of KpFtsZtr–Mb in the *P*2_1_2_1_2_1_ space group, Mb-free KpFtsZtr (PDB code:6LL5), and FtsZ from *Pseudomonas aeruginosa* (PDB code:2VAW) are superimposed. 2*F*_o_–*F*_c_ map around GDP is contoured at 1.0σ for the KpFtsZtr–Mb. **c**, Close-up view around the FtsZ–Mb interface in the KpFtsZtr–Mb complex in the *P*2_1_2_1_2_1_ space group. Mb is colored in gray. **d**, Crystal packing of KpFtsZtr–Mb in the *P*2_1_ space group. A pair of three FtsZ monomers in the asymmetric unit is colored by temperature factor. Mbs are colored in gray. The inset shows a close-up view around GDP in the upper two molecules.

Next, we investigated whether the binding of Mb(S1) affected the GTPase activity of KpFtsZ. The GTPase activity assay was performed using the enzyme coupling method^33^, where GTP hydrolysis is coupled with a decrease in absorbance at 340 nm. As KpFtsZ supplemented with the Mb showed slightly decreased but comparable activity to KpFtsZ alone (Fig. 1f), the binding of Mb had little effect on the GTPase activity of KpFtsZ. We also observed *E. coli* cells overproducing the Mb by fluorescence microscopy to evaluate the effects of Mb on the localization and formation of Z-rings. Mbs are cysteine-free and are therefore functional in a reducing environment inside the cells. When EGFP was fused at the C-terminus end of the Mb, the Mb-EGFP-expressing cells showed an elongated morphology compared to cells expressing EGFP alone (Fig. 1g), indicating that the Mb inhibits cell division. The Mb-EGFP dots were observed every 2–5 μm, suggesting that Mb-EGFP was bound to FtsZ protofilaments forming Z-rings. Thus, it is likely that the Mb does not affect the formation of the endogenic Z-ring but inhibits its constriction.

### Crystal structures of the KpFtsZ–Mb and EcFtsZ–Mb complexes

Two crystal forms of KpFtsZtr complexed with the Mb and two crystal forms of truncated EcFtsZ (residues 11–316; EcFtsZtr) complexed with the Mb were obtained under the same crystallization conditions. The structures of the two crystal forms (*P*2_1_2_1_2_1_ and *P*2_1_) of the KpFtsZtr–Mb and EcFtsZtr–Mb complex were determined using molecular replacement with KpFtsZtr (Protein Data Bank (PDB) code:6LL5^23^) at resolutions of 2.20 and 2.50 (KpFtsZtr–Mb) and 1.84 and 2.60 Å (EcFtsZtr–Mb), respectively (Table 1). The *P*2_1_2_1_2_1_ and *P*2_1_ crystals of the KpFtsZtr–Mb and EcFtsZtr–Mb complexes showed the R conformation and contained one and three complex molecules, respectively, per asymmetric unit. As the *P*2_1_2_1_2_1_ and *P*2_1_ crystal structures of KpFtsZtr–Mb were mostly similar to those of EcFtsZtr–Mb (a root-mean-square deviation (r.m.s.d.) of 0.187 Å for 280 C_α_ atoms for the *P*2_1_2_1_2_1_ datasets and 0.326 Å for 286 C_α_ atoms for the *P*2_1_ datasets; Extended Data Fig. 3), we hereafter describe the crystal structures of KpFtsZtr– Mb.

**Table 1.** Crystallographic data collection statistics of KpFtsZtr–Mb and EcFtsZtr–Mb complexes.

| Dataset | KpFtsZtr–Mb |  | EcFtsZtr–Mb |  |
| --- | --- | --- | --- | --- |
| PDB accession code | 8GZV | 8GZW | 8GZX | 8GZY |
| X-ray source | SPring-8 | SPring-8 | SPring-8 | SPring-8 |
|  | BL44XU | BL44XU | BL44XU | BL44XU |
| Wavelength (Å) | 0.900 | 0.900 | 0.900 | 0.900 |
| Temperature (°C) | –173 | –173 | –173 | –173 |
| Resolution range | 35.54–2.20 | 40.05–2.50 | 35.2–1.84 | 35.25–2.60 |
|  | (2.28–2.20) | (2.59–2.50) | (1.91–1.84) | (2.69–2.60) |
| Space group | *P*2<sub>1</sub>2<sub>1</sub>2<sub>1</sub> | *P*2<sub>1</sub> | *P*2<sub>1</sub>2<sub>1</sub>2<sub>1</sub> | *P*2<sub>1</sub> |
| *Unit-cell parameters* |  |  |  |  |
| *a* (Å) | 45.29 | 88.47 | 45.11 | 89.34 |
| *b* (Å) | 64.56 | 66.93 | 63.30 | 66.23 |
| *c* (Å) | 124.65 | 102.17 | 123.05 | 102.79 |
| $\alpha$ (°) | 90.00 | 90.00 | 90.00 | 90.00 |
| $\beta$ (°) | 90.00 | 92.06 | 90.00 | 92.67 |
| $\gamma$ (°) | 90.00 | 90.00 | 90.00 | 90.00 |
| Total reflections | 250,724 (23,545) | 287,239 (30,389) | 412,374 (39,436) | 253,395 (26,839) |
| Unique reflections | 19,223 (1,877) | 41,534 (4,148) | 31,362 (3,066) | 37,170 (3,686) |
| Redundancy | 13.0 (12.5) | 6.9 (7.3) | 13.1 (12.9) | 6.8 (7.3) |
| Completeness (%) | 99.86 (99.73) | 99.69 (99.86) | 99.95 (100.0) | 99.6 (99.8) |
| $R_{\text{merge}}$ | 0.09836 (0.9052) | 0.07902 (0.7059) | 0.09167 (1.621) | 0.05329 (0.6137) |
| $I/\sigma$ | 17.67 (4.06) | 15.57 (2.62) | 19.50 (1.77) | 19.95 (2.96) |
| $CC\ 1/2$ | 0.999 (0.938) | 0.998 (0.866) | 0.999 (0.661) | 0.999 (0.925) |
| *Refinement* |  |  |  |  |
| $R_{\text{work}}/R_{\text{free}}$ | 0.205/0.264 | 0.215/0.284 | 0.205/0.243 | 0.214/0.296 |
| *No. of atoms* |  |  |  |  |
| Protein | 2,897 | 8,684 | 2,870 | 8,607 |
| Ligand | 28 | 84 | 28 | 84 |
| Water | 46 | 40 | 123 | 9 |
| *R. m. s. deviation* |  |  |  |  |
| Bond length (Å) | 0.008 | 0.008 | 0.007 | 0.009 |
| Bond angles (°) | 0.99 | 1.08 | 1.05 | 1.12 |

Statistics for the highest-resolution shell are shown in parentheses.

**Fig. 3.**
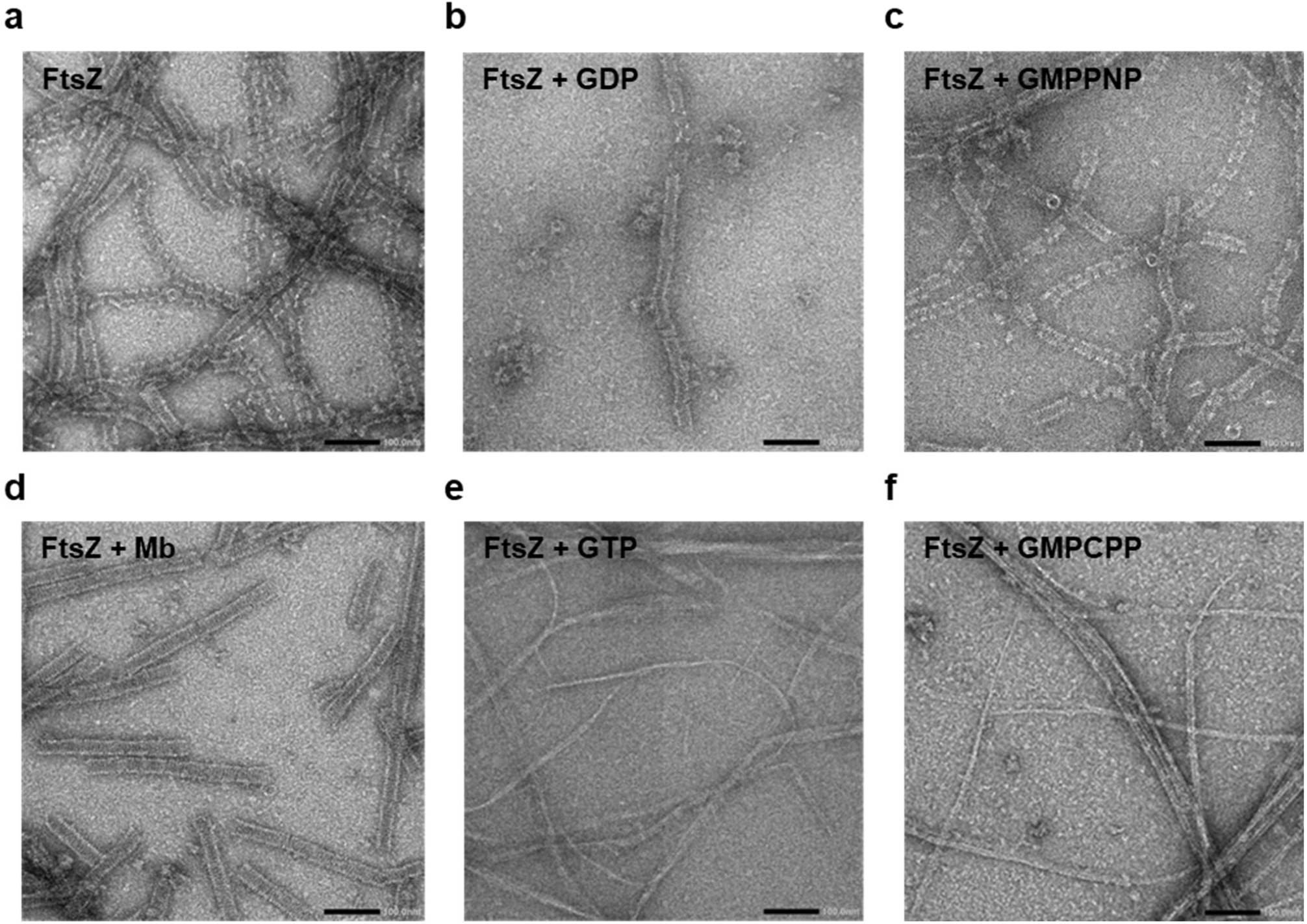
Various morphologies of KpFtsZ filaments. **a**, Typical negative stain micrograph of 1.0 mg ml^−1^ FtsZ. **b–f**, Typical negative stain micrographs of 1.0 mg ml^−1^ FtsZ supplemented with (**b**) 1 mM GDP, (**c**) 1 mM GMPPNP, (**d**) 1.2 molar excess of Mb, (**e**) 1 mM GTP, and (**f**) 1 mM GMPCPP. Scale bar represents 100 nm.

The monomeric structures of Mb-bound KpFtsZtr in the two space groups were similar (r.m.s.d. of 0.454 Å for 265 C_α_ atoms; Fig. 2a). In contrast, the monomeric structures of Mb-bound KpFtsZtr in the two space groups and Mb-free KpFtsZtr (PDB code:6LL5^23^) were relatively different (r.m.s.d. of 1.005 Å for 272 C_α_ atoms for the *P*2_1_2_1_2_1_ dataset and 1.080 Å for 284 C_α_ atoms for the *P*2_1_ dataset). Large structural differences between these structures were observed in the T3 loop and KpFtsZtr–Mb interface region. The structure of the T3 loop in KpFtsZtr–Mb showed a closed conformation similar to that of FtsZ from *Pseudomonas aeruginosa* (Fig. 2b), whereas the T3 loop in Mb-free KpFtsZtr adopted a unique open conformation due to the crystal packing^23^. In the KpFtsZtr–Mb complex, the Mb interacted with the central helix (H7 helix), the H6-H7 loop, and a β-sheet in the C-terminal domain. The interaction between FtsZ and the Mb was mainly mediated by Arg residues in the FG loop of the Mb (Figs. 1b and 2c; Extended Data Fig. 4). The binding region of the Mb was distant from the GTP-binding site of FtsZ, which is consistent with the finding that the GTPase activity of KpFtsZ was not affected by Mb binding.

**Fig. 4.**
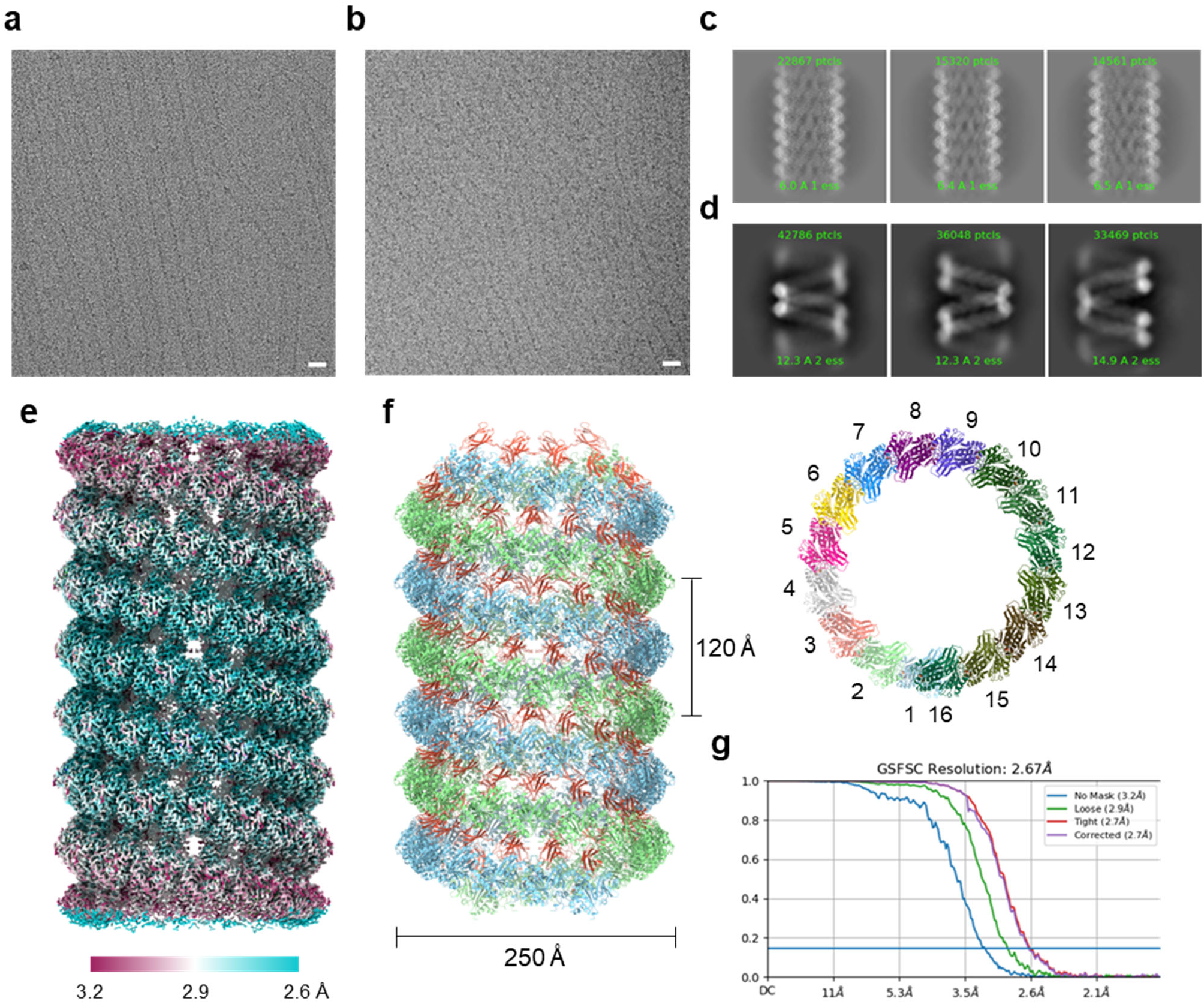
CryoEM analysis of KpFtsZ–Mb double helical tube. **a**,**b**, Typical raw micrographs (x 600,000) of (**a**) 4.0 mg ml^−1^ FtsZ–Mb and (**b**) 4.0 mg ml^−1^ FtsZ. Scale bar represents 20 nm. **c**,**d**, Part of selected 2D class averages of (**c**) FtsZ–Mb and (**d**) FtsZ datasets aligned in the descending order of particle numbers from left to right and top to bottom. **e**, Final sharpened map of FtsZ–Mb double helical tube. The local resolution distributions are colored as in the color bar. **f**, Fitted model of a FtsZ filament in two orthogonal views; side view, left panel; and top view, right panel. In the left panel, two FtsZ protofilaments are shown in cyan and green. Mb molecules are shown in red. In the right panel, each FtsZ–Mb complex is shown in different colors. **g**, The FSC curve for the final map of FtsZ–Mb dataset. The horizontal blue line indicates the FSC = 0.143 criterion.

The *P*2_1_2_1_2_1_ crystal structures of KpFtsZtr–Mb showed straight protofilaments in the crystal (Extended Data Fig. 5). The molecular architecture of the protofilaments was very similar to that of KpFtsZ (PDB code:6LL5^23^). In contrast, the *P*2_1_ KpFtsZtr–Mb structures demonstrated that the three KpFtsZtr molecules in the asymmetric unit form a curved protofilament. The superposition of the C_α_ atoms between the *P*2_1_2_1_2_1_ and *P*2_1_ KpFtsZtr protofilaments composed of three KpFtsZtr molecules indicated that the protofilaments of *P*2_1_ KpFtsZtr are curved. The GDP portion of the middle and bottom KpFtsZtr molecules was well ordered and exhibited relatively low temperature factors (the average C_α_ temperature factor is 49.9 and 45.3 Å^2^ for GDP), compared to the average temperature factor of 80.4 Å^2^ for the GDP of the top KpFtsZtr molecule (Fig. 2d). The crystal packing revealed a tight packing of trimers but fewer interactions around the GDP of the top molecules.

**Fig. 5.**
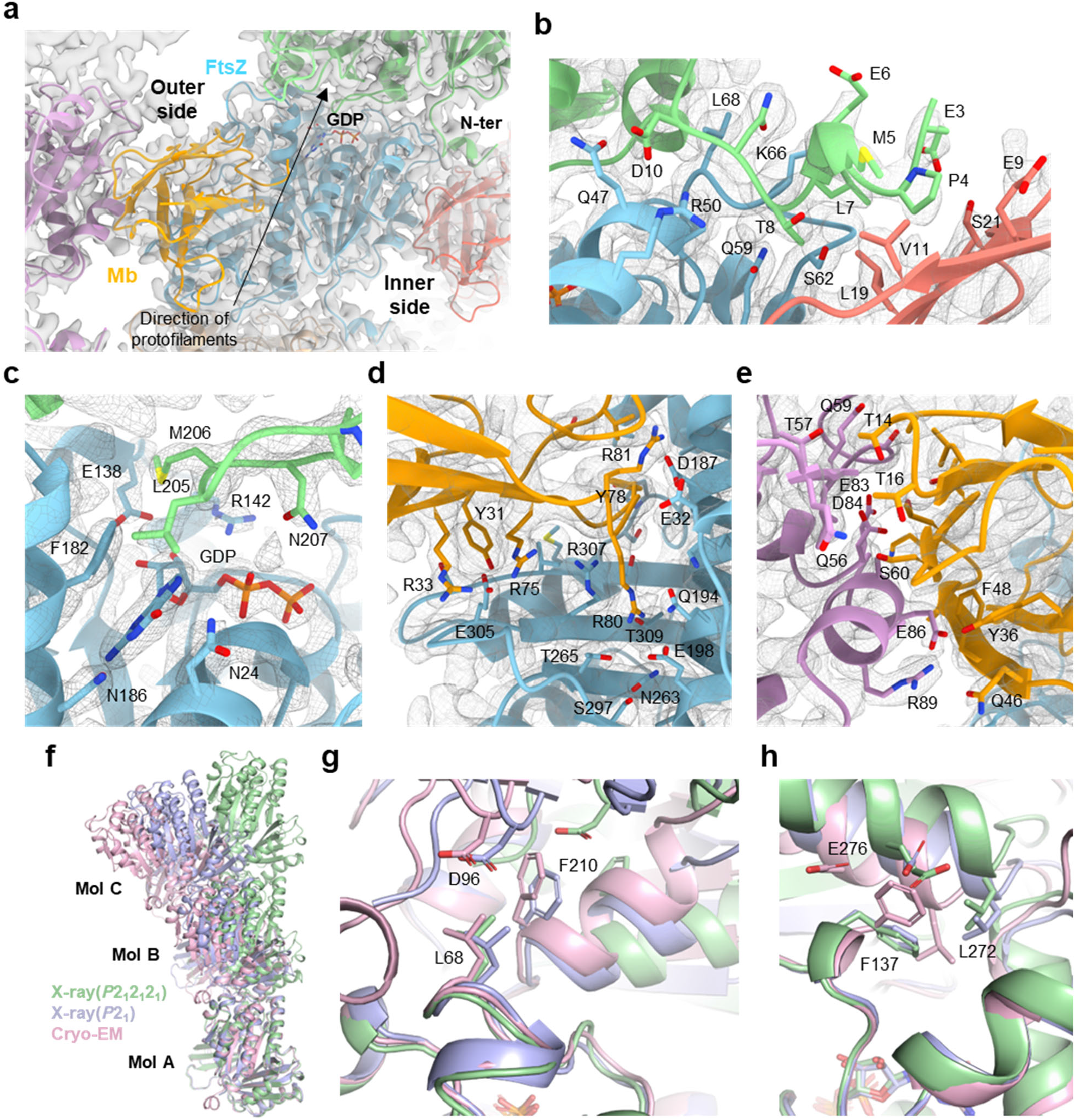
Interactions in the KpFtsZ–Mb double helical tubes. **a**, Close-up view around an FtsZ molecule (shown in cyan) and Mb (show in yellow) fitted into the final sharpened map. The molecules contributing to longitudinal interaction (in a protofilament) are shown in green and wheat. The molecules participating in Mb-mediated lateral interaction (between protofilaments) are shown in pink and red. **b–e**, Close-up view around (**b**) N-terminal interface, (**c**) GDP interface, (**d**) FtsZ–Mb interface within a complex, and (**e**) FtsZ–Mb interface between two complexes. Residues located in the surfaces are shown in stick. The coloring is the same as in (**a**), and the map is shown in mesh. **f**, Structure comparison of KpFtsZ protofilaments of the three datasets. **g**,**h**, Close-up views of interfaces between mol A and B in (**f**). (**g**) is the N-terminal side and (**h**) is the Mb side.

### Observation of various types of KpFtsZ filaments

To investigate the effects of nucleotides and the Mb on filament formation, we observed KpFtsZ filaments by negative staining EM under various conditions. KpFtsZ alone at a concentration of 1.0 mg ml^−1^ showed thick and flexible tubes, as observed previously^27–30^ (Fig. 3a). The addition of GDP and GMPPNP did not significantly affect the tube shape (Fig. 3b,c). These tubes appeared fragile and many nicks were found. Unexpectedly, supplementation with the Mb significantly changed the shape of the tubes to thicker and straight (Fig. 3d). These tubes appeared to have a periodic structure, and much fewer nicks were observed. The addition of GTP and GMPCPP also dramatically altered the properties of the FtsZ filaments. Very thin and bundled filaments were observed under these conditions (Fig. 3e,f), as observed in several previous studies. Most filaments were straight, but they seemed to have some flexibility. We excluded the GTP and GMPCPP conditions from the specimen preparation for cryoEM structure analysis because of the thinness of the filaments.

### CryoEM of KpFtsZ–Mb double helical filaments

We prepared two specimens for cryoEM: one with 4.0 mg ml^−1^ KpFtsZ supplemented with 1 mM GMPPNP; and the other additionally supplemented with 1.2× molar excess of the Mb. Similar to the observation by negative staining EM, the filaments of KpFtsZ–Mb seemed to be straighter and more rigid than those of KpFtsZ lacking the Mb (Fig. 4a,b). After data collection and image processing, the 2D class averages of KpFtsZ–Mb showed periodic and helical structures (Fig. 4c). In contrast, the 2D class averages of KpFtsZ showed slightly different spring-like structures (Fig. 4d), and we could not reconstruct a high-resolution 3D map probably because of the high flexibility of the filaments. After refining the helical parameters, the reconstituted 3D map of KpFtsZ–Mb represented a double helical filament with a diameter of 250 Å, a helical rise of 7.70 Å, and a helical twist of –23.40° (Fig. 4e). The C2 symmetry was imposed as the two KpFtsZ protofilaments formed double helices, and the Mb molecules were located between the two protofilaments (Fig. 4f, left panel). We constructed a model of the KpFtsZ–Mb complex and found that each protofilament contained 15.4 molecules per turn (Fig. 4f, right panel). The resolution of the final reconstructed 3D map was 2.67 Å (Fig. 4g; Extended Data Fig. 6). Although the densities corresponding to a C-terminus region composed of residues 317–383 was not observed, the visible C-terminus of FtsZ (residue 316) was located outside the tube (Extended Data Fig. 7), as previously observed^27^.

**Fig. 6.**
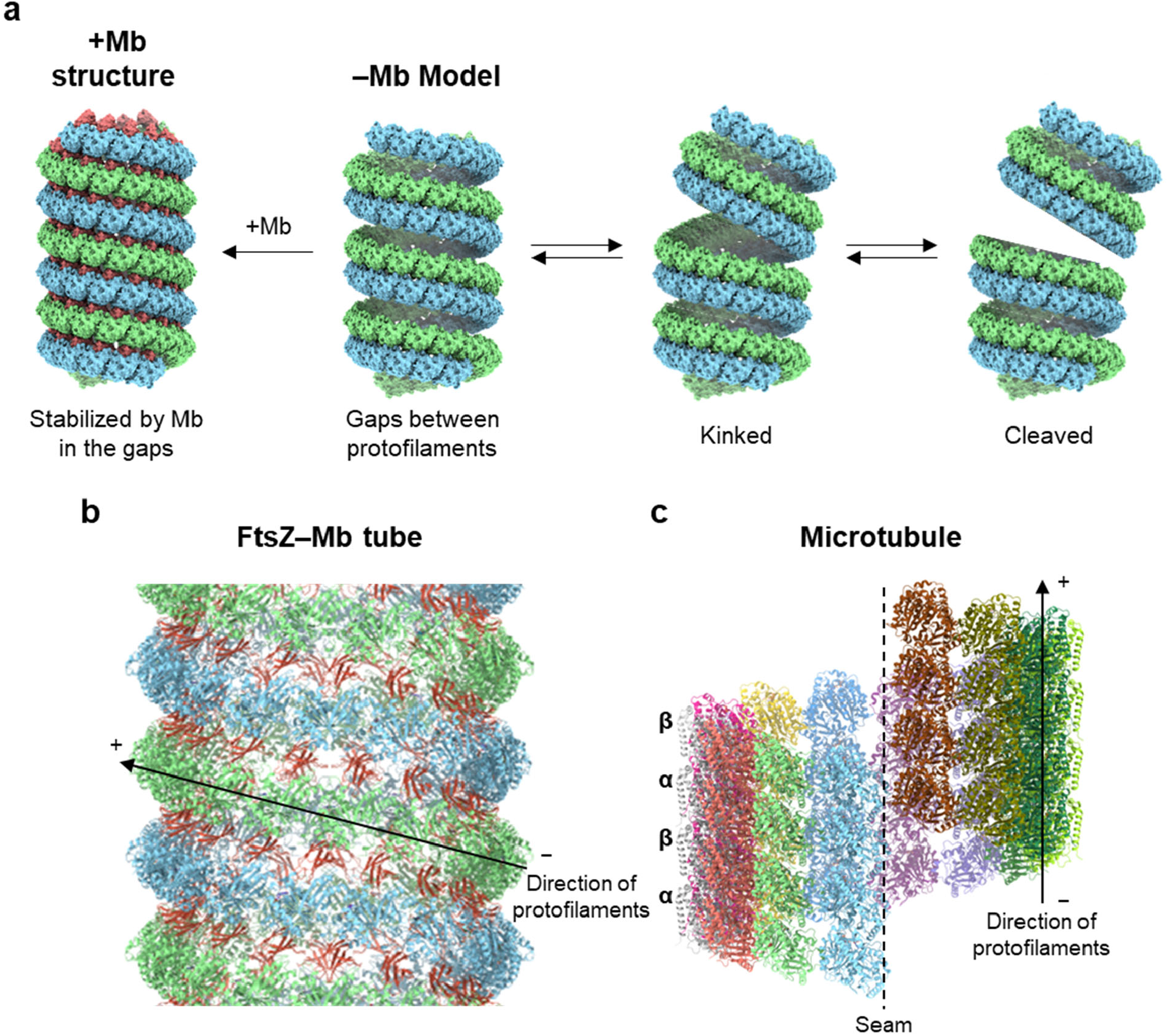
Dynamics and structure comparison of the FtsZ tubes. **a**, Dynamics model of FtsZ double helical tubes. **b**, Side view of KpFtsZ–Mb tube. The coloring is the same as in Fig. 4f. The direction of the protofilaments are indicated by an arrow with plus (+) and minus (−) end. **c**, Side view of a microtubule from *Sus scrofa* (PDB code:6DPU). The α- and β-tubulin molecules within a single protofilament are shown in the same colors.

We built a model of the KpFtsZ–Mb complex and found that the overall structure showed the R conformation; it was very similar to the crystal structures (r.m.s.d. = 0.387 Å over 366 C_α_ atoms for the *P*2_1_ dataset and 0.568 Å over 360 C_α_ atoms for the *P*2_1_2_1_2_1_ dataset) (Extended Data Fig. 7), and the longitudinal GDP-mediated interaction within the protofilament was maintained (Fig. 5a, see also Supplementary Video 1). In contrast to most previous FtsZ structures, the N-terminal region composed of residues 3–10 was ordered and clearly visible (Fig. 5b). Residues 4–8 formed a short 3_10_ helix stabilized by the N-terminal cap of Pro4, which was located in a hydrophobic pocket formed by Glu9, Val11, Leu19, and Ser21 of the adjacent Mb. We found GDP in the binding pocket instead of GMPPNP (Fig. 5c). GDP was sandwiched by two KpFtsZ molecules in a protofilament, similarly to many previously reported crystal structures. A cluster of salt bridges and hydrophilic interactions between KpFtsZ and the Mb within the complex were maintained (Fig. 5d). On the other hand, the Mb contributed to lateral hydrophilic interactions with the neighboring protofilament (Fig. 5e).

To evaluate the curvature of the FtsZ protofilaments, we superimposed the three models of KpFtsZ–Mb trimers generated from the two crystal structures and one cryoEM structure with the first molecule (mol A) and found that the cryoEM protofilament showed a larger curvature than the others (Fig. 5f). Focusing on the N-terminal side of the interface between mol A and mol B, Leu68, Asp96, and Phe210 formed hydrophobic interactions in the curved protofilaments (cryoEM and *P*2_1_ crystal structures), whereas Asp96 and Phe210 were dissociated with a large rearrangement of the N-terminal domain of mol B in the straight protofilament (*P*2_1_2_1_2_1_ crystal structure, Fig. 5g). On the opposite Mb side, Phe137 was flipped to allow the large movement of Leu272 and Glu276 on the α11 helix in the cryoEM structure (Fig. 5h). The interface areas between the KpFtsZ monomers in the cryoEM protofilament (1065 Å^2^ and 1068 Å^2^) were larger than those of the crystal structures (703 Å^2^ for *P*2_1_2_1_2_1_ and 931 Å^2^ and 822 Å^2^ for *P*2_1_, Table 3).

**Table 2.** CryoEM data collection and processing.

|  |  |
| --- | --- |
| Dataset | KpFtsZ–Mb |
| EMDB accession code | EMD-34429 |
| PDB accession code | 8H1O |
| **Data collection and processing** |  |
| Magnification | 60,000 |
| Voltage (kV) | 300 |
| Electron exposure ($e^-/\text{\AA}^2$) | 60 |
| Defocus range ($\mu\text{m}$) | −0.5 to −2.0 |
| Pixel size ($\text{\AA}$) | 0.883 |
| Symmetry imposed | C2 helical |
| Imported movies (no.) | 3,096 |
| Initial particle images (no.) | 181,247 |
| Final particle images (no.) | 90,711 |
| Map resolution ($\text{\AA}$) | 2.67 |
| FSC threshold | 0.143 |
| Map sharpening B factor ($\text{\AA}^2$) | −81.3 |
| Helical parameters |  |
| Rise ($\text{\AA}$) | 7.70 |
| Twist ($^\circ$) | −23.40 |
| **Refinement** |  |
| Initial model used (PDB code) | 8GZV |
| Model resolution ($\text{\AA}$) | 2.6/2.6/2.8 |
| FSC threshold | 0/0.143/0.5 |
| Model vs. Data CC (mask) | 0.87 |
| (volume) | 0.82 |
| Model composition |  |
| Non-hydrogen atoms | 2975 |
| Protein residues | 404 |
| Ligands | 1 (GDP) |
| R.m.s. deviations |  |
| Bond lengths ($\text{\AA}$) | 0.003 |
| Bond angles ($^\circ$) | 0.532 |
| Validation |  |
| MolProbity score | 1.27 |
| Clashscore | 5.05 |
| Poor rotamers (%) | 0 |
| Ramachandran plot |  |
| Favored (%) | 98.8 |
| Allowed (%) | 1.2 |
| Disallowed (%) | 0 |

**Table 3.**
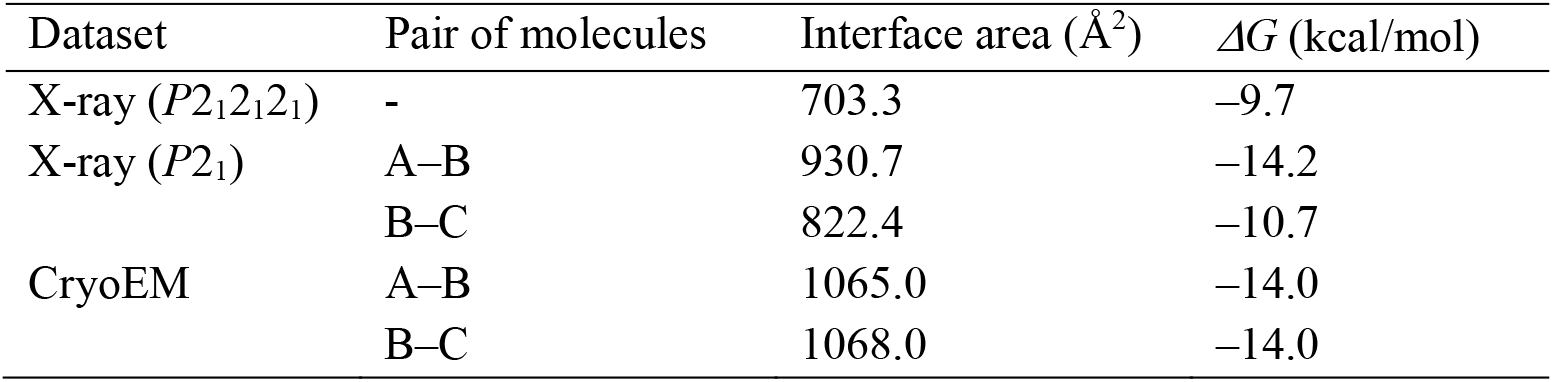
Interface areas between two KpFtsZ molecules in a protofilament.

### Discussion

Based on the unrestricted binding property of FtsZ to multiple nucleotide-bound states, together with its non-impeditive molecular size (∼10 kDa), we envisioned that Mbs would be beneficial for the structural analysis of various types of FtsZ assemblies. Although the double helical tubes of FtsZ that we determined were stabilized by an Mb, tubes with similar diameters were also observed without the Mb. These tubes appeared much shorter and more flexible than those with the Mb (Fig. 4b). The two protofilaments in the KpFtsZ–Mb tube aligned at regular intervals due to the stabilization by the Mb (Fig. 4c), whereas the two protofilaments in the tube without the Mb were arranged such that they form large gaps (Fig. 4d). These gaps allow the tubes to be bent or kinked like a spring, or even cleaved into short tubes, which could provide a flexibility to the FtsZ tubes (Fig. 6a).

The model of the double helical FtsZ tube was proposed in 1999 using electron micrographs and their optical diffraction patterns^34^. Helical tubes were also observed in EcFtsZ, but not in MtbFtsZ (Fig. S14 of Ref. 29). They were observed only in the presence of GDP, and the L272E and A181E mutations inhibited the formation of these tubes. These two residues were located on the Mb side interface in the protofilament (Fig. 5h), which suggests the formation of protofilaments with similar interfaces in the previous helical tubes. Another study also reported the helical tubes of MjFtsZ (Fig. 2d of ^30^). These results inspired us to hypothesize that the large helical tubes of FtsZ are not limited to a few species but reflect the structure of the Z-ring in some phases during cell division. Twisted helical filaments of FtsZ were observed in the presence of ZapA (Fig. 1b and 6b of ^31^). ZapA is a crosslinker for FtsZ protofilaments and increases the spatial order of the filament network without affecting the treadmilling^35^. ZapA or other accessory proteins may function as Mb to regulate the lateral interactions between the FtsZ protofilaments. Different sizes of FtsZ crosslinkers could result in different spacings between the protofilaments at different stages of Z-ring constriction. Additionally, the helical FtsZ tube seems to allow membrane anchor proteins such as FtsA and ZipA to bind, as the visible C-terminus of FtsZ (residue 316) is located outside the tube (Extended Data Fig. 7). This result is consistent with previous observations for FtsZ with C-terminal tags^27^. Notably, ZipA and FtsA* (a gain-of-function mutant of FtsA) can stabilize highly curved protofilaments and minirings of FtsZ^9^.

From our observations, the addition of GTP and GMPCPP dramatically changed the KpFtsZ filament properties to those of single straight filaments, as observed frequently, but GDP and GMPPNP did not (Fig. 3). This means that the shapes of the KpFtsZ filaments are not affected by the presence of γ-phosphate (GTP-like or GDP-like). One possible explanation is that GMPPNP could not bind KpFtsZ because of its low affinity^1^, which is supported by the fact that our cryoEM structure of the FtsZ helical tube contains only GDP, despite the addition of 1 mM GMPPNP. As GMPCPP is known as a slowly hydrolyzable analog, GTP hydrolysis would be required for the formation of long and straight FtsZ protofilaments.

FtsZ is a bacterial tubulin homologue. Although there have been a large number of structural studies on microtubules, those on FtsZ have been limited to protofilament units. The tubes with a larger order than the protofilaments have been observed in previous EM studies as described above. The question of whether these tubes contain protofilament-like interactions, and if so, which direction the protofilaments go, had remained to be elucidated, because of the lack of high-resolution structure of the FtsZ tubes in solution. In this study, we found that KpFtsZ can also form microtubule-like tubes with a similar diameter of 250 Å, and the high-resolution cryoEM structure of the tube revealed detailed molecular interactions. The largest difference between the FtsZ tubes and microtubules was the direction of the protofilaments. Note that the intermolecular interactions within a protofilament are mediated by GDP or GTP. In the FtsZ tube, the protofilaments grew diagonally to the tube axis (Fig. 6b), whereas dimers of α- and β-tubulin aligned along the microtubule axis to form protofilaments^36^ (Fig. 6c). The FtsZ tube contains only a single unit and does not have a seam, but the microtubules consist of two units and have a seam. The monomeric structure of KpFtsZ in the helical tube is not very similar to those of tubulin monomers (r.m.s.d. = 2.820 over 101 C_α_ atoms for α-tubulin and 4.655 over 161 C_α_ atoms for β-tubulin, PDB code:6DPV^36^, Extended Data Fig. 8), and the interface areas between tubulin monomers in a protofilament (∼1726 Å^2^ for α–β interface and ∼1702 Å^2^ for β–α interface) are much larger than those between FtsZ monomers in a protofilament in the double helical tube (∼1065 Å^2^, Table 3). This is because the C-terminal helical region of tubulin participates in the interactions between α- and β-tubulin (Extended Data Fig. 9). The corresponding region of FtsZ is a flexible linker that is disordered. The smaller number of interactions between FtsZ monomers may affect the dynamics of protofilaments, such as the directions of curvatures^27^ and lead to different properties of tubes compared to microtubules (Extended Data Fig. 10). Additionally, many microtubule accessory proteins are known to help regulate microtubule formation. Some FtsZ accessory proteins may also work in a similar way to regulate the shape of the Z-ring in some steps during cell division.

As cell division proceeds, FtsZ needs to form Z-rings with various diameters and shapes. A large ring diameter indicates almost straight protofilaments, and a constricted ring should consist of highly curved protofilaments, as proposed more than 20 years ago^37^. The FtsZ protofilaments in the previous crystal structures are straight or weakly bent and are relatively stable. In contrast, the FtsZ protofilaments in the helical tube are the most largely curved and are unstable without spacer molecules such as Mb preventing the tubes from dissociation. It is still unclear whether a structure like the double helical FtsZ tubes actually exists in cells because the artifactual Mb and a graphene support layer stabilized the FtsZ double helical tube in this study. However, similar highly curved protofilaments would be necessary in a constricted Z-ring and stabilized by FtsZ accessory proteins *in vivo*. Structural flexibility or various curvatures of the FtsZ protofilaments must be regulated by many accessory proteins and are important for cell division.

### Online content

Any methods, additional references, Nature Research reporting summaries, source data, extended data, supplementary information, acknowledgements, peer review information; details of author contributions and competing interests; and statements of data and code availability are available at …

## Online Methods

### Protein expression and purification

We used full-length and truncated versions of KpFtsZ and EcFtsZ in this study. The FtsZ derivatives were prepared as a fusion protein C-terminal to His_6_ and a TEV cleavage site or another fusion protein C-terminal to His_6_, a biotin-acceptor tag (Avi-tag) and a TEV cleavage site using modified pCold I vector (Takara Bio Inc.). These proteins were produced and purified as previously described^1^, except that the biotinylated proteins were produced in BL21(DE3) cells containing the pBirAcm plasmid (Avidity) in the presence of 50 μM D-biotin for *in vivo* biotinylation. A monobody binding to the two FtsZs was prepared as a His_10_-tagged protein using the pHFT2 vector as previously described^2,3^. *E. coli* derived proteins, including the maltose binding protein (MBP), the phosphoenolpyruvate carboxylase (EcPEPC), the ribonuclease H2 (EcRNH2), and a mutant of the adenylate kinase (EcAdktm; A55C/C77S/V169C)^4^ were also prepared as N-terminally His_6_-tagged and biotinylated proteins using the pHBT vector^5^ to validate off-target binding of monobodies. These off-target proteins were produced with the same protocol for the biotinylated FtsZ proteins. The FtsZ derivatives were purified using a 5 ml HisTrap HP column (Cytiva) and a 5 ml HiTrap Q HP column (Cytiva), and their monodispersity was confirmed by a Superdex200 size-exclusion column (Cytiva). The monobody and off-target proteins were purified using a 5 ml HisTrap HP column and further purified with a ProteoSEC Dynamic 16/60 3-70 HR size-exclusion column (Protein Ark) or a Superdex200 size-exclusion column. For enzyme assay, crystallization and cryoEM experiments, the affinity tag for each protein was removed using TEV protease.

### Monobody generation

The monobody libraries used and general selection methods have been described previously^6,7^. The buffers used for binding reaction and washing were TBSB (20 mM Tris-HCl pH 7.4, containing 150 mM NaCl and 1 mg ml^−1^ bovine serum albumin) and TBSBT (TBSB and 0.05% (v/v) Tween 20), respectively, for phage display selection experiments. Four rounds of phage display selection against EcFtsZ were performed. The EcFtsZ concentrations used for rounds 1–2 and 3–4 were 100 and 50 nM, respectively. Monobody-displayed phages were captured onto a biotinylated target immobilized to streptavidin-coated magnetic beads (Z5481/2, Promega) and then eluted in 0.1 M Gly-HCl, pH 2.1.

After four rounds of selection, individual clones from the enriched library were isolated on agar plates and six of them were subjected to phage ELISA to validate the binding and specificity against EcFtsZ and KpFtsZ. For phage preparation and ELISA assay, procedures were performed essentially as described previously^8^. Five clones binding to both EcFtsZ and KpFtsZ were selected, and their amino acid sequences were deduced by DNA sequencing.

Affinity of the generated monobody was determined using yeast-surface display as described previously^7^, except that the buffers used for binding reaction and washing were HBSB (20 mM HEPES-NaOH pH 7.5, containing 150 mM NaCl and 1 mg ml^−1^ bovine serum albumin) and HBSBT (HBSB and 0.05% (v/v) Tween 20), respectively. Yeast cells displaying a monobody were incubated with varying concentrations of EcFtsZ or KpFtsZ, washed with the buffer and stained with appropriate fluorescently labeled secondary detection reagents, prior to analysis on a Muse flow cytometer (Millipore). *K*_d_ values were determined from plots of the median fluorescent intensity against FtsZ concentration by fitting the 1:1 binding model using SigmaPlot software (Systat Software). Experiments were performed in triplicate. The detail of the gating strategy is described in Supplementary Fig. 1.

### Effect of nucleotides on monobody binding

Effect of nucleotides on monobody binding was examined using yeast-surface display as described above, except that an appropriate concentration of EcFtsZ or KpFtsZ was pre-incubated with or without a nucleotide/nucleotide-analogue (1 mM for GDP, GTP, and GMPPNP and 0.1 mM for GMPCPP) in HBSB for 30 min on ice, and then the mixture was transferred to the wells where monobody binding took plate. The concentrations of EcFtsZ and KpFtsZ used were 1.5 μM and 10 μM, respectively, to account for different affinity. Experiments were performed in triplicate.

### Gel filtration chromatography of KpFtsZtr–Mb complex

Gel filtration chromatography of KpFtsZtr–Mb complex was carried out using a 24 ml Superdex 200 column (Cytiva) equilibrated with 20 mM HEPES-NaOH pH 7.5, 150 mM NaCl. 18.9 mg ml^−1^ KpFtsZtr and 9.9 mg ml^−1^ Mb were mixed and incubated at room temperature for 1 hr before being subjected to gel filtration chromatography. The isolated complex had an apparent molecular mass of 49.9 kDa as estimated from gel filtration chromatography.

### GTPase activity assay

GTPase assay of KpFtsZ was performed using a coupled enzyme assay^9^. In this method, a GTP hydrolysis is coupled to NADH oxidation with pyruvate kinase (PK) and lactate dehydrogenase (LDH). NADH absorbance decrease at 340 nm was measured, which is proportional to the rate of GTP hydrolysis. In our experiments, the sequential reactions were performed in 50 mM Tris-HCl pH 7.5, 5 mM MgCl_2_, 200 mM KCl, 50 U ml^−1^ PK, 50 U ml^−1^ LDH, 1 mM phosphoenolpyruvate (PEP), 1 mM GTP, 0.2 mM NADH, and 0.81 mg ml^−1^ FtsZ at 37 °C. Total reaction volume was 1 ml. GTPase activity was calculated from the slope of the fitted line in the area with linearly decreasing absorbance at 340 nm divided by the molar extinction coefficient of NADH (6220 M^−1^cm^−1^), FtsZ molar concentration (20 μM), and the path length (1 cm). Data are obtained as the mean of three independent experiments with standard deviation.

### Fluorescence microscopy

For imaging experiments, *E. coli* BL21(DE3) was transformed with Mb-EGFP or EGFP expressing vector and grown in LB medium until OD_600_ = 0.388. The proteins were induced by the addition of 0.5 mM isopropyl β-_D_-thiogalactopyranoside (IPTG). 2 μl of the samples were applied to the glass slide after 25 min of induction and then covered with 18 mm × 18 mm cover glass. Cells were imaged using inverted fluorescence microscopy (Nikon, Eclipse Ti2) equipped with a 100× oil immersion objective lens (Nikon). The microscope system was operated using the NIS-elements (Nikon).

#### Protein crystallization

Purified KpFtsZtr and EcFtsZtr at a concentration of 10 mg ml^−1^ was mixed with 1.2× molar excess of Mb. The final concentration of KpFtsZtr and EcFtsZtr, and Mb are 8 mg ml^−1^ and 0.4 mg ml^−1^, respectively. KpFtsZtr–Mb and EcFtsZtr-Mb crystals belonging to the space group *P*2_1_ and *P*2_1_2_1_2_1_ were obtained by hanging-drop vapour diffusion at 293 K (1 μl protein solution + 1 μl reservoir solution) with the same reservoir solution consisting of 0.2 M sodium formate, 0.1 M Bis-Tris propane pH6.5, and 20% PEG3350.

### Crystallographic data collection, processing, and refinement

Crystals were flash-cooled in a stream of nitrogen at 100 K without cryoprotectants after mounting in a loop. X-ray diffraction data were collected at a wavelength of 0.900 Å on the micro-focus beamline BL44XU at SPring-8, Hyogo, Japan using an EIGER X 16M detector (Dectris). The datasets for both KpFtsZtr–Mb and EcFtsZtr–Mb were integrated, and scaled using the KAMO system^10^ which runs BLEND^11^, XDS, and XSCALE ver. 5^12^ (February 2021) automatically. The phases for each structure were determined by molecular replacement with MOLREP^13^ in the CCP4 suite ver. 7.1^14^ using the previously determined structure of KpFtsZtr (PDB code:6LL5)^1^ as the search model. Each model was refined with REFMAC ver. 5.8.0267^15^ and PHENIX ver. 1.19.1^16^, with manual modification using Coot ver. 0.8.6^17^. The refined structures were validated with MolProbity^18^. Data-collection and refinement statistics are shown in Table 1.

### Negative staining

One side of amorphous carbon grids were hydrophilized by glow discharge using a JEC-3000FC sputter coater (JEOL). 3 μl of purified KpFtsZ at a concentration of 1.0 mg ml^−1^ was loaded on the grid and blotted. Then the grid was immediately stained with 3 μl of 2% uranyl acetate solution and blotted, and this process was repeated three times. The grid was air-dried for 30 min. Specimens of KpFtsZ supplemented with any of 1 mM GDP, GMPPNP, GTP, GMPCPP, or 1.2× molar excess of Mb were prepared in the same way. Images were taken using JEM-1400Flash (JEOL, Japan) operated at 100 kV.

### CryoEM specimen preparation and data collection

Purified KpFtsZ at a concentration of 4.0 mg ml^−1^ was mixed with 1 mM GMPPNP, 0.12 mM PC190723, and 1.2× molar excess of Mb, and the solution was incubated on ice for 20 min. Another KpFtsZ alone sample without adding PC190723 and Mb was also prepared in the same way. Epoxidized graphene grid (EG-grid)^19^ was used to stabilize and increase the number of FtsZ filaments. 3 μl of the 0.01 M NaOH and 1% (v/v) epichlorohydrin water solution was applied to the ClO_2_^•^-oxidized graphene grids to prepare EG-grids. Then 3 μl of the protein solution was applied to the EG-grid. After incubation at room temperature for 5 min, the grids were blotted with a force of 0 and a time of 3 sec (KpFtsZ–Mb complex) or with a force of –10 and a time of 1.5 sec (KpFtsZ) in a Vitrobot Mark IV chamber (Thermo Fisher Scientific, USA) equilibrated at 4 °C and 100% humidity and then immediately plunged into liquid ethane. Excessive ethane was removed with filter paper, and the grids were stored in liquid nitrogen. CryoEM image datasets were acquired using SerialEM ver. 3.8^20^ and JEM-Z300FSC (CRYO ARM™ 300: JEOL, Japan) operated at 300 kV with a K3 direct electron detector (Gatan, Inc.) in CDS mode. The Ω-type in-column energy filter was operated with a slit width of 20 eV for zero-loss imaging. The nominal magnification was 60,000×, corresponding to 0.88 Å per pixel. Defocus varied between –0.5 μm and –2.0 μm. Each movie was fractionated into 60 frames (0.0505 s each, total exposure: 3.04 s) with a total dose of 60 e^−^/Å^2^.

### CryoEM image processing and model building

The images were processed using cryoSPARC ver. 3.3.1^21^. 3,096 movies of KpFtsZ–Mb dataset were imported and motion corrected, and the contrast transfer functions (CTFs) were estimated. 2,889 micrographs whose CTF max resolutions were beyond 5 Å were selected. 181,247 particles were automatically picked using filament tracer job with the parameters; filament diameter, 250 Å; separation distance between segments, 0.5; minimum filament length to consider, 3. At an early stage of processing, particles were extracted with the box size of 480 pixel with 2x binning. After several rounds of 2D classification, helix refine, and symmetry search, sub-optimal values of helical rise and twist were found. Then particles were extracted with the box size of 600 pixel without binning. After two rounds of 2D classification, 90,711 particles were selected. The particles were subjected to helix refine job with the initial helical parameters and without imposing symmetry. After global and local CTF refine jobs, another helix refine job was performed. Then the final helix refine job was run with C2 symmetry. The refined values of helical rise and twist were 7.70 Å and –23.40°, respectively, and the final map resolution (FSC=0.143) reached 2.67 Å. The dataset of KpFtsZ was also processed in the similar way, but the helical parameters were difficult to refine due to the structural flexibility of the filaments, and the map resolution was limited to 7–8 Å.

The monomeric model of KpFtsZ–Mb was built using the *P*2_1_2_1_2_1_ crystal structure of KpFtsZtr– Mb complex as an initial model. After the initial model was manually fitted into the map using UCSF Chimera ver. 1.15^22^ and modified in Coot ver. 0.9.6^23^, real-space refinement was performed in PHENIX ver. 1.19.2^16^. The model was validated using MolProbity^18^, and this cycle was repeated several times. The whole tube model (containing 100 KpFtsZ–Mb complexes) was generated using the following command in UCSF Chimera: “sym #1 group C2*H,7.703,-23.398,50,-25 center 264.9,264.9,264.9”. Interface areas were calculated with PISA server^24^. Figures were prepared using UCSF Chimera ver. 1.15^22^, ChimeraX ver. 1.1^25^, PyMOL ver. 2.5.0 (Schrödinger, LLC, USA), and LigPlot^+^ ver. 2.2.4^26^. The parameters are summarized in Table 2.

### Reporting summary

Further information on research design is available in the Nature Research Reporting Summary linked to this article.

## Data availability

Coordinates and structure factors have been deposited in the Protein Data Bank (PDB) under the accession numbers 8GZV for KpFtsZtr–Mb (*P*2_1_2_1_2_1_), 8GZW for KpFtsZtr–Mb (*P*2_1_), 8GZX for EcFtsZtr–Mb (*P*2_1_2_1_2_1_), and 8GZY for EcFtsZtr–Mb (*P*2_1_). CryoEM map and atomic coordinate have been deposited in the Electron Microscopy Data Bank (EMDB) and the PDB under the accession codes EMD-34429 and 8H1O, respectively. All other data are available from the authors upon reasonable request.

## Acknowledgements

This work was supported by: JSPS KAKENHI grant JP18K05445 (S-i.T.), JP21K05386 (S-i.T.), JP18K06094 (H.M.), JP19H04735 (H.M.), and JP20K22630 (J.F.); the Sasakawa Scientific Research Grant from the Japan Science Society 2018-3011 (S-i.T.), and 2022-4052 (H.A.); JST OPERA (Open Innovation with Enterprises, Research Institute and Academia) grant JPMJOP1861 (T.I., K.N.); AMED BINDS (Platform Project for Supporting Drug Discovery and Life Science Research (BINDS) grant JP21am0101117 and JP22ama121003 (K.N.); AMED CiCLE (Cyclic Innovation for Clinical Empowerment) grant JP17pc0101020 (K.N.); JEOL YOKOGUSHI Research Alliance Laboratories of Osaka University (K.N.); the Cooperative Research Program of Institute for Protein Research, Osaka University (CR-20-02 and CR-21-02). We thank Kiyoaki Arakawa, Miho Emori, Takahiro Hayashi, Momoka Iiyama, and Naoki Okazaki for help in monobody screening by phage display, and Yoshie Kushima for help in negative staining. This work has been performed under the approval of the SPring-8 Program Advisory Committee (Proposal Nos. 2020A2536, 2020A2544, 2020A6557, 2021A6623, 2021A6648, and 2021B1002).

## Author contributions

J.F., S-i.T., and H.M. designed the research. H.A., M.H., and S-i.T. acquired the monobody and analyzed its binding affinity. T.Y. N.K., N.Kamimura, and H.M. prepared the protein sample, performed activity assay, binding assay, crystallization, crystallographic data collection, processing, refinement. T.Y. and N.Kamimura observed *E. coli* cells using fluorescent microscopy. J.F., N.K., N.Kamimura, and H.M. performed negative staining, and cryoEM data collection. J.F. performed cryoEM image processing and model building. J.F., H.A, T.Y., S-i.T., and H.M. prepared the figures and wrote the first draft of the manuscript. T.I. and K.N. helped to analyze and interpret the data, and critically revise the manuscript. J.F., S-i.T. and H.M. conceptualized the study, developed the study design, supervised the authors throughout the study, and provided expertise in manuscript preparation. All authors read and approved the reviewed manuscript.

## Additional information

**Extended data** is available for this paper at https://doi.org/

**Supplementary information** The online version contains supplementary material available at https://doi.org/

**Correspondence and requests for materials** should be addressed to Shun-ichi Tanaka or Hiroyoshi Matsumura.

### Peer review information

**Reprints and permissions information** is available at http://www.nature.com/reprints

**Extended Data Fig. 1.**
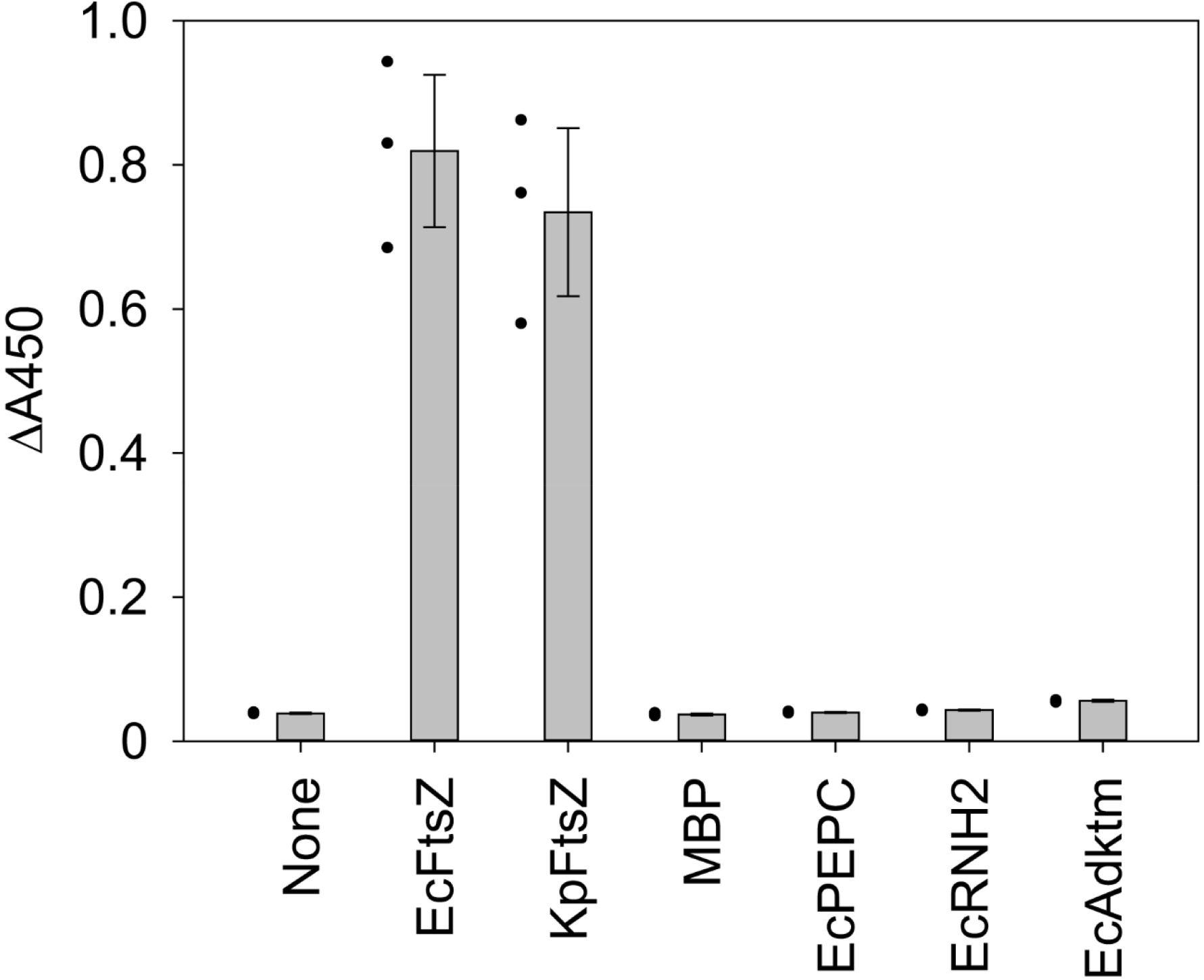
Phage ELISA analysis of binding of Mb(S1) to its cognate and off-target proteins. The binding of Mb(S1)-phages to KpFtsZ- or EcFtsZ-coated wells as a cognate target and MBP-, EcPEPC-, EcRNH2-, or EcAdktm-coated wells as an off-target was measured. The error bars indicate s.d. from three independent experiments, and where none are visible, the errors are within the size of the markers.

**Extended Data Fig. 2.**
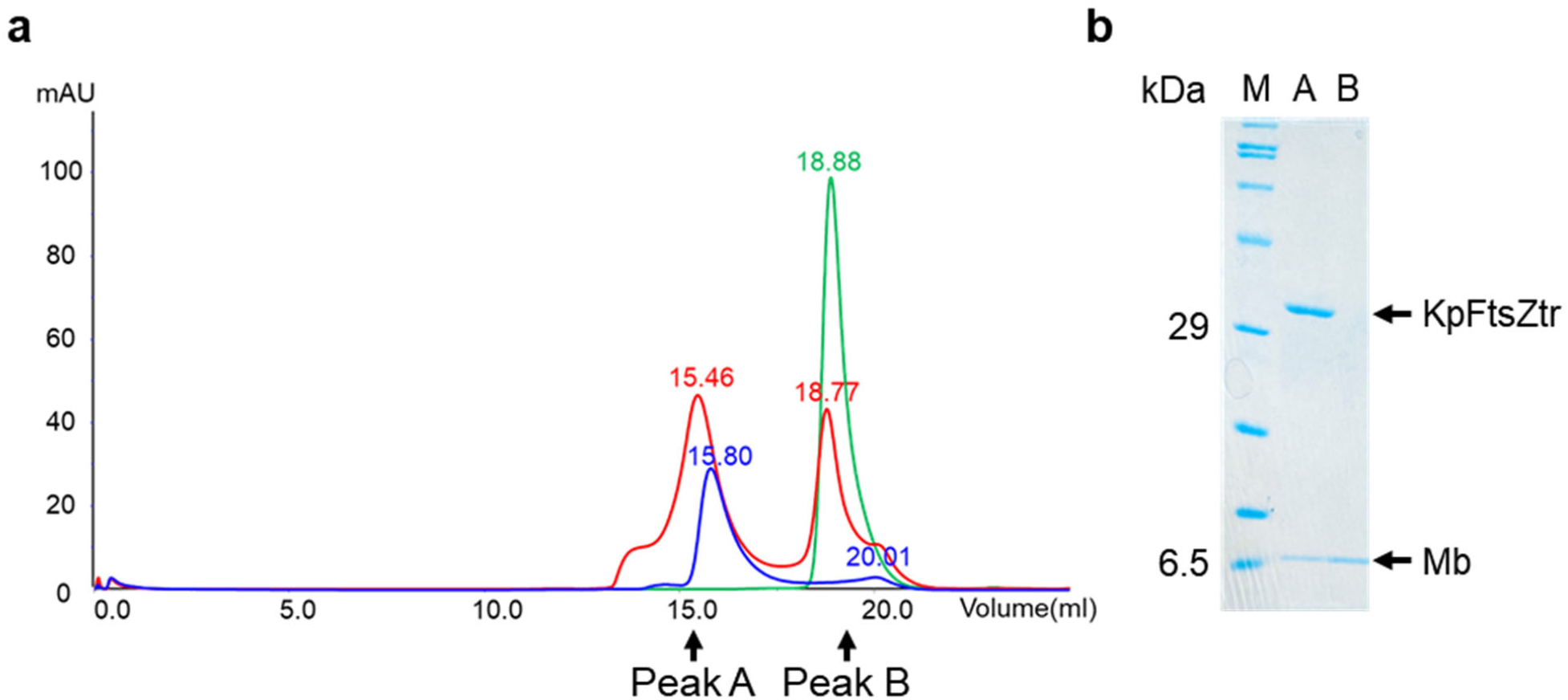
Formation of KpFtsZtr–Mb complex. **a**, Gel filtration chromatograms of KpFtsZtr (blue curve), Mb (green curve), and the KpFtsZtr–Mb complex (red curve). **b**, SDS-PAGE analysis of the different fractions of the KpFtsZtr–Mb complex analysis. ‘M’ denotes protein marker; ‘A’ denotes peak A fraction; ‘B’ denotes peak B fraction.

**Extended Data Fig. 3.**
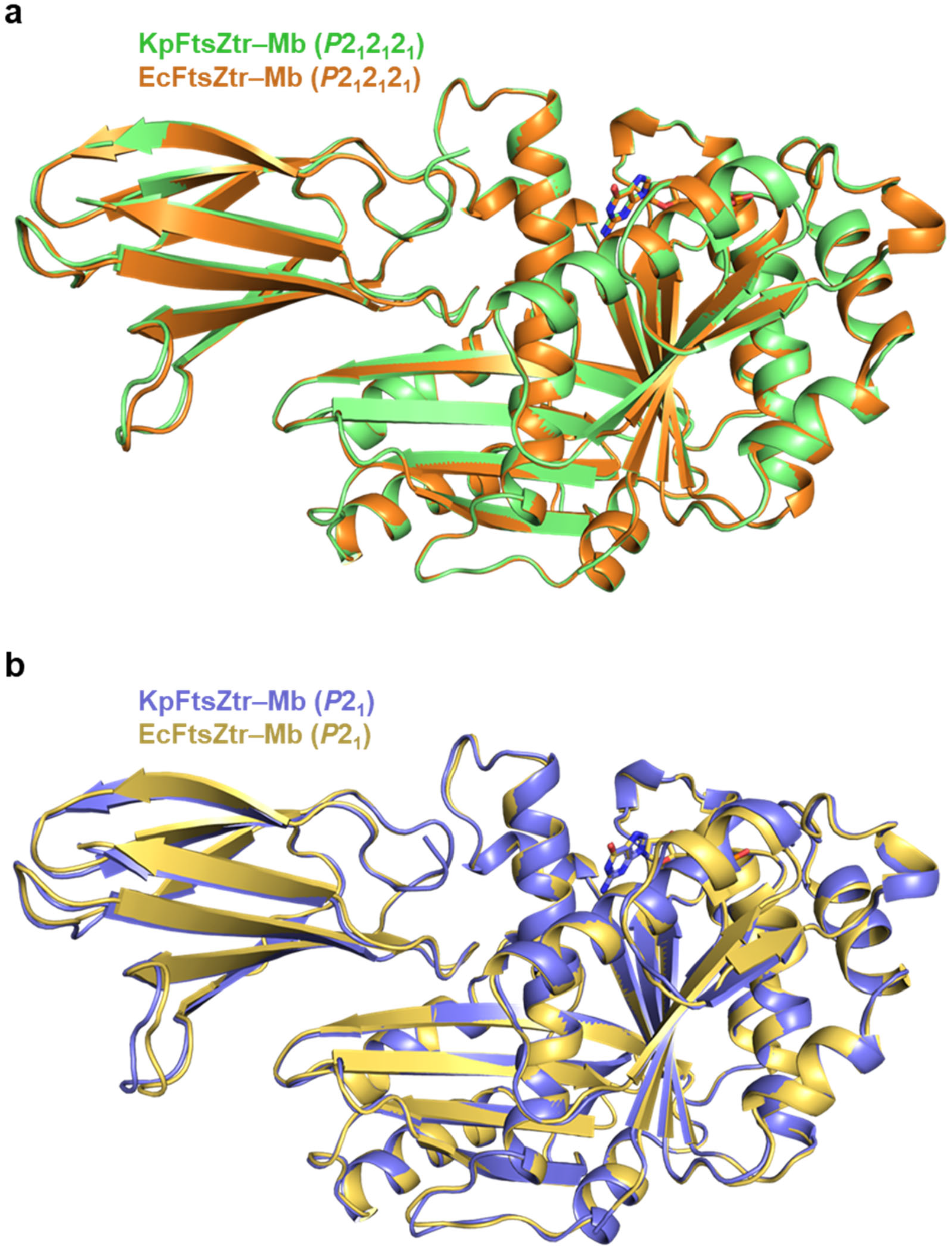
Structure comparison of KpFtsZtr–Mb and EcFtsZtr–Mb. **a**,**b**, Superimposed crystal structures of the KpFtsZtr–Mb with EcFtsZtr–Mb (**a**) in the *P*2_1_2_1_2_1_ space group and (**b**) in the *P*2_1_ space group.

**Extended Data Fig. 4.**
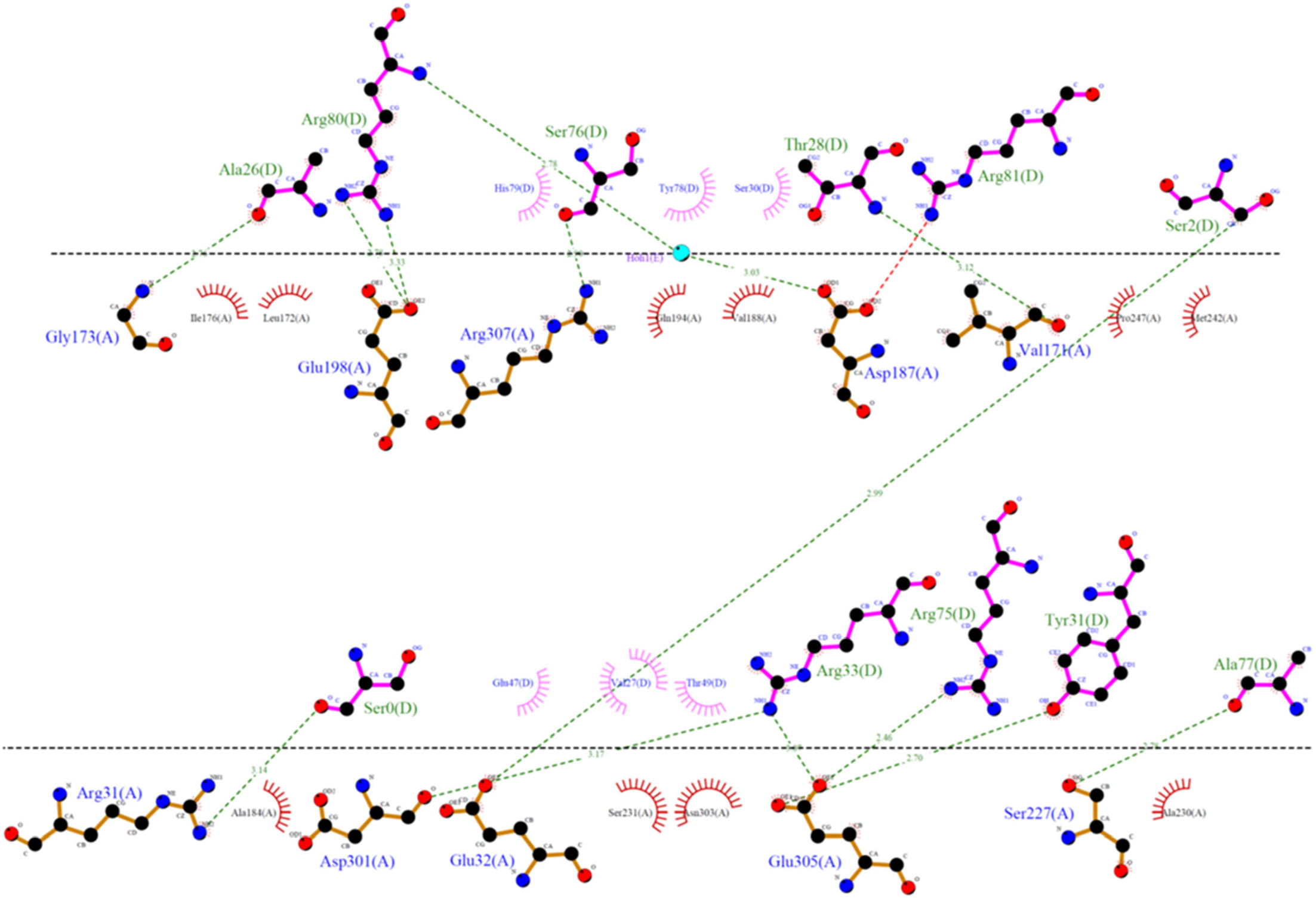
Detailed interfacial interactions between KpFtsZ and Mb in the *P*2_1_2_1_2_1_ dataset. Interactions were detected and drawn with LigPlot^+^.

**Extended Data Fig. 5.**
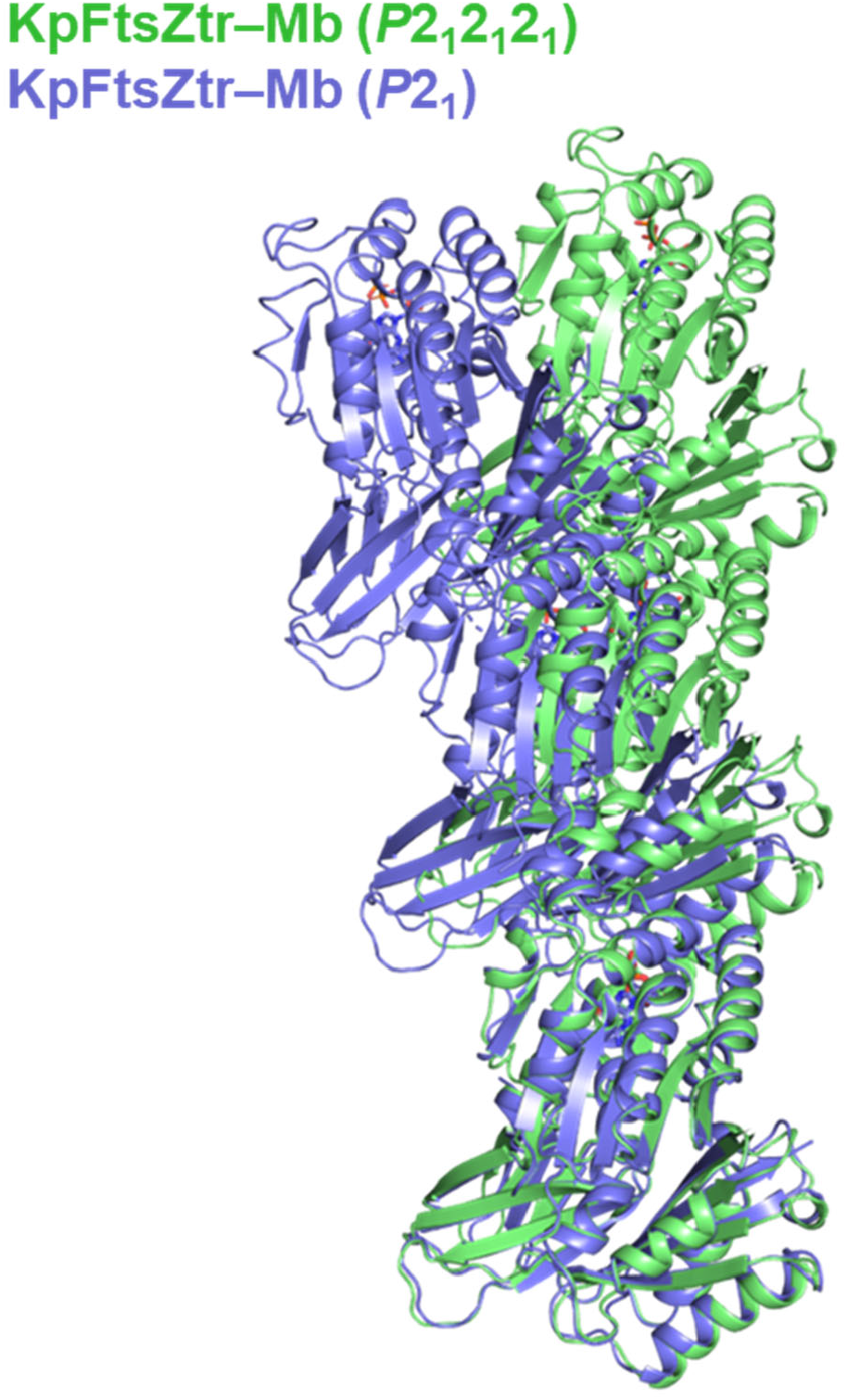
Structure comparison of KpFtsZtr–Mb protofilaments in the two different space groups.

**Extended Data Fig. 6.**
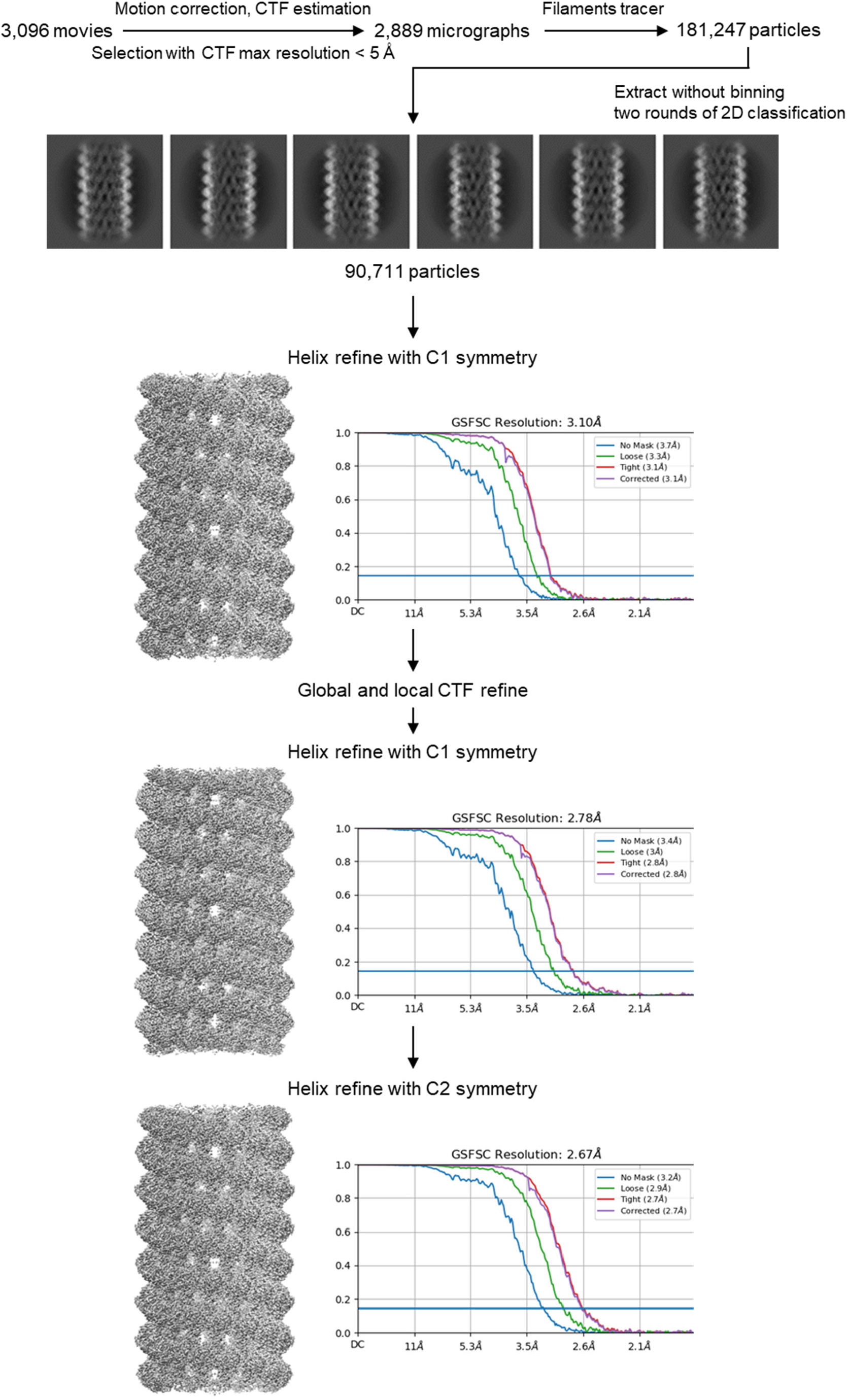
CryoEM data processing workflow of KpFtsZ–Mb complex.

**Extended Data Fig. 7.**
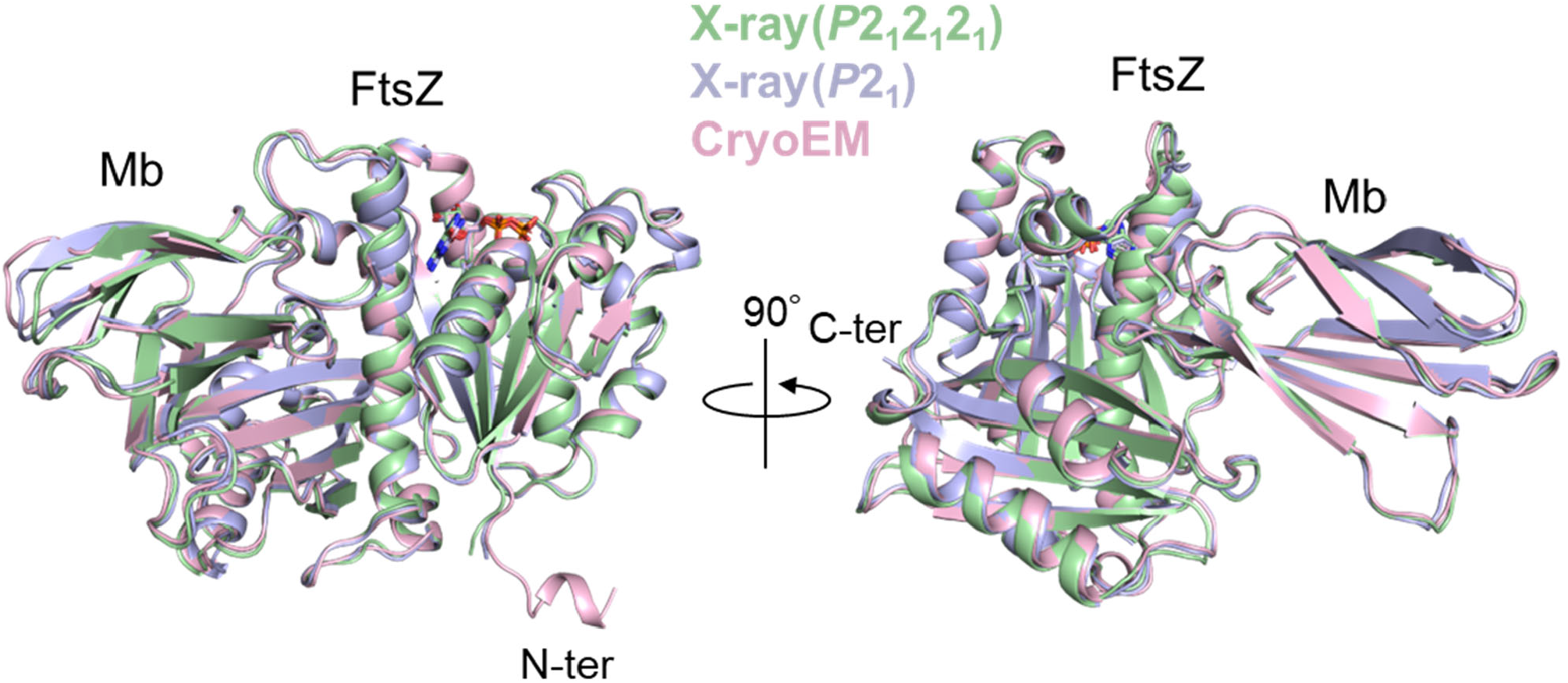
Structure comparison of KpFtsZ monomers in the crystal and CryoEM structures. Two orthogonal views are shown in the left and right panels.

**Extended Data Fig. 8.**
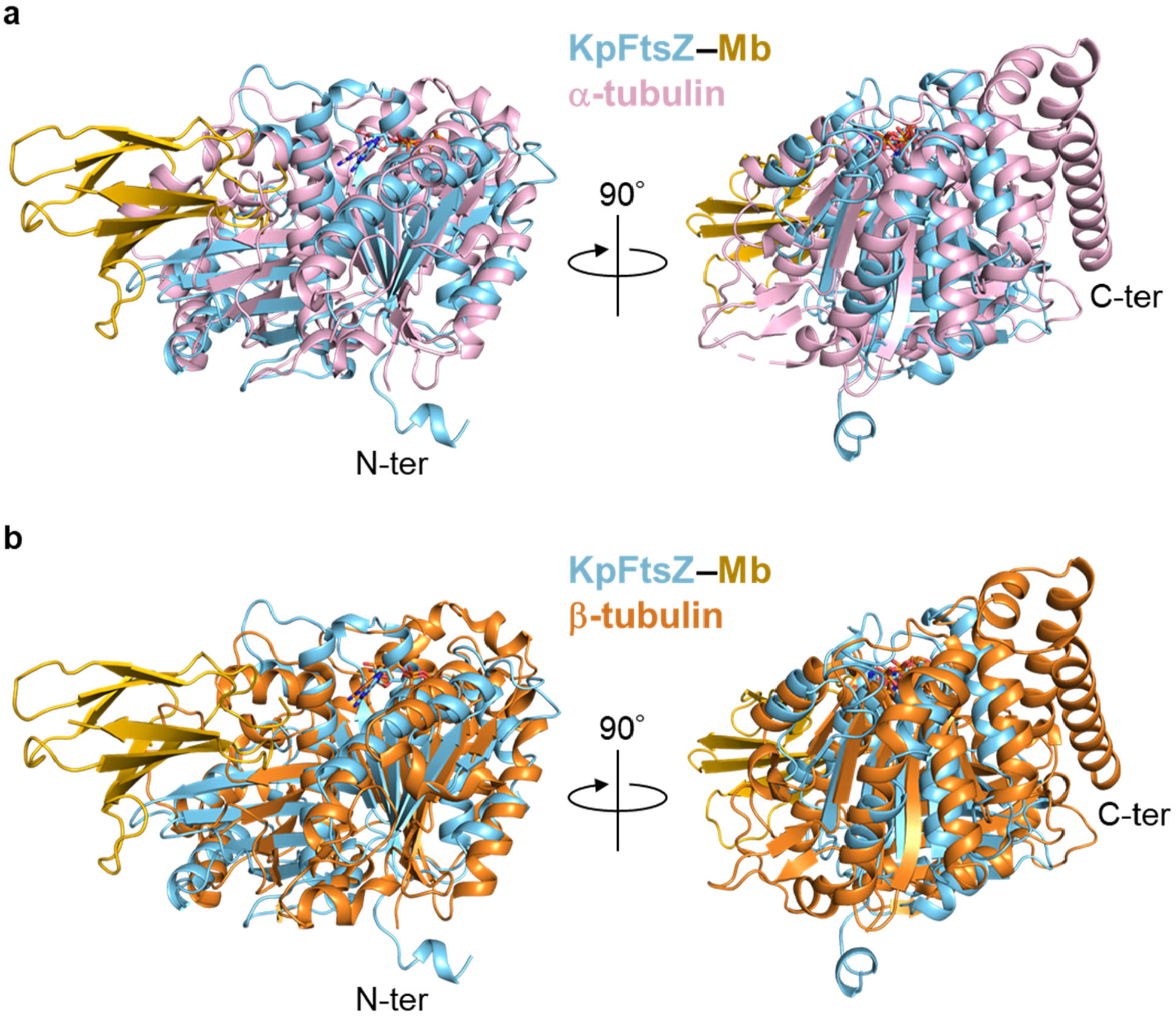
Structure comparison between KpFtsZ and tubulin monomers. **a**,**b**, Superimposed structures of the KpFtsZ–Mb complex in the cryoEM structure with (**a**) a-tubulin and (**b**) b-tubulin (PDB code:6DPV). Two orthogonal views are shown in the left and right panels. Only FtsZ moiety is used for superimposition for the KpFtsZ–Mb complex.

**Extended Data Fig. 9.**
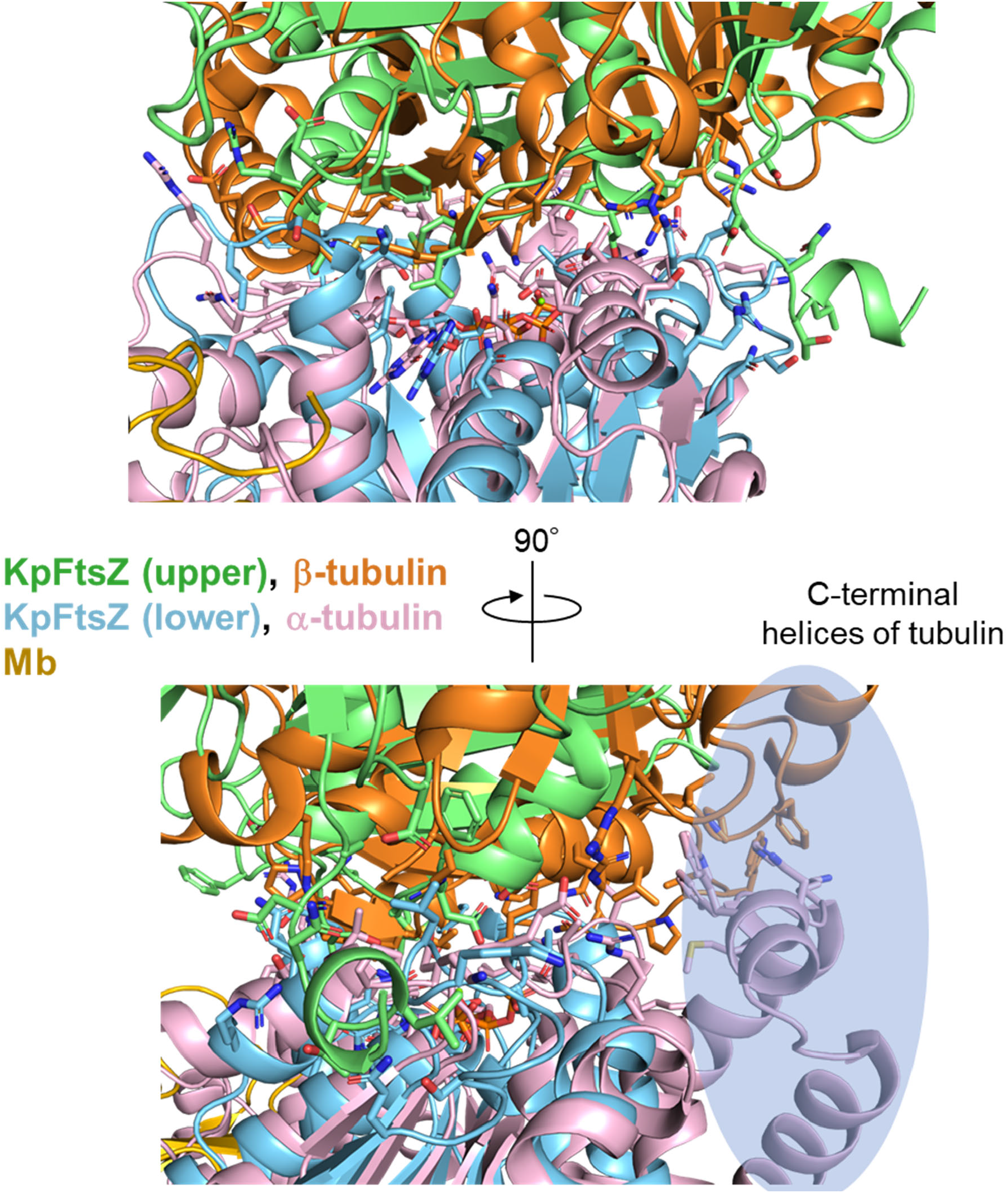
Interfacial interactions between the KpFtsZ protofilament and α-β tubulin dimers. The structures of lower KpFtsZ (cyan) and a-tubulin (pink) are superimposed. Two orthogonal views are shown in the upper and lower panels. C-terminal helices region of tubulin is highlighted in the lower panel.

**Extended Data Fig. 10.**
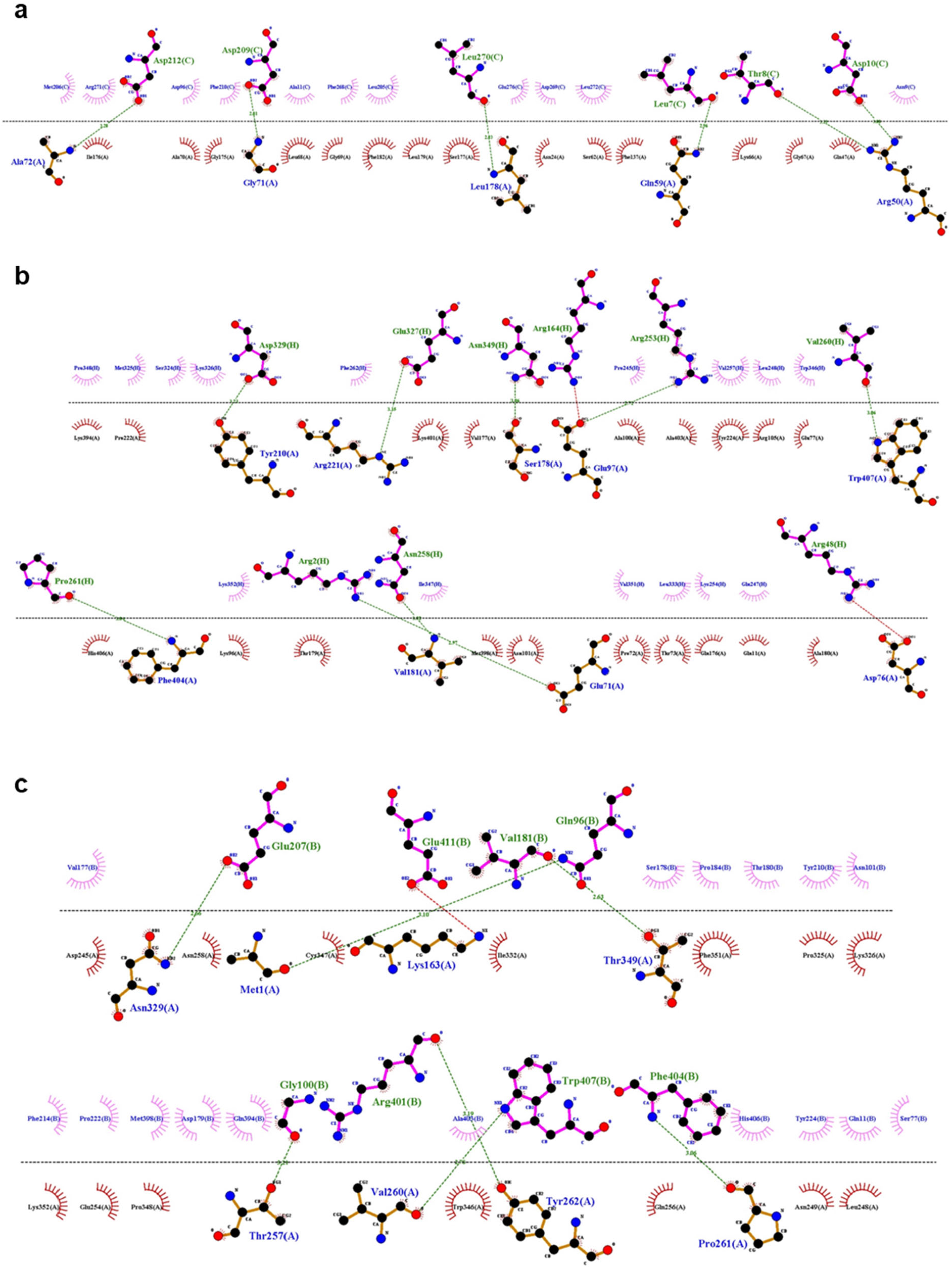
Detailed interfacial interactions between monomers in the KpFtsZ and tubulin protofilaments. **a–c**, Interactions in (**a**) KpFtsZ interface, (**b**) α (chain A)–β (chain H) tubulin interface, and (**c**) β (chain B)–α (chain A) tubulin interface (PDB code:6DPV). Interactions were detected and drawn with LigPlot^+^.

