## Supplementary Information for "High-resolution structure of a microtubule-like tube composed of FtsZ–monobody complexes"

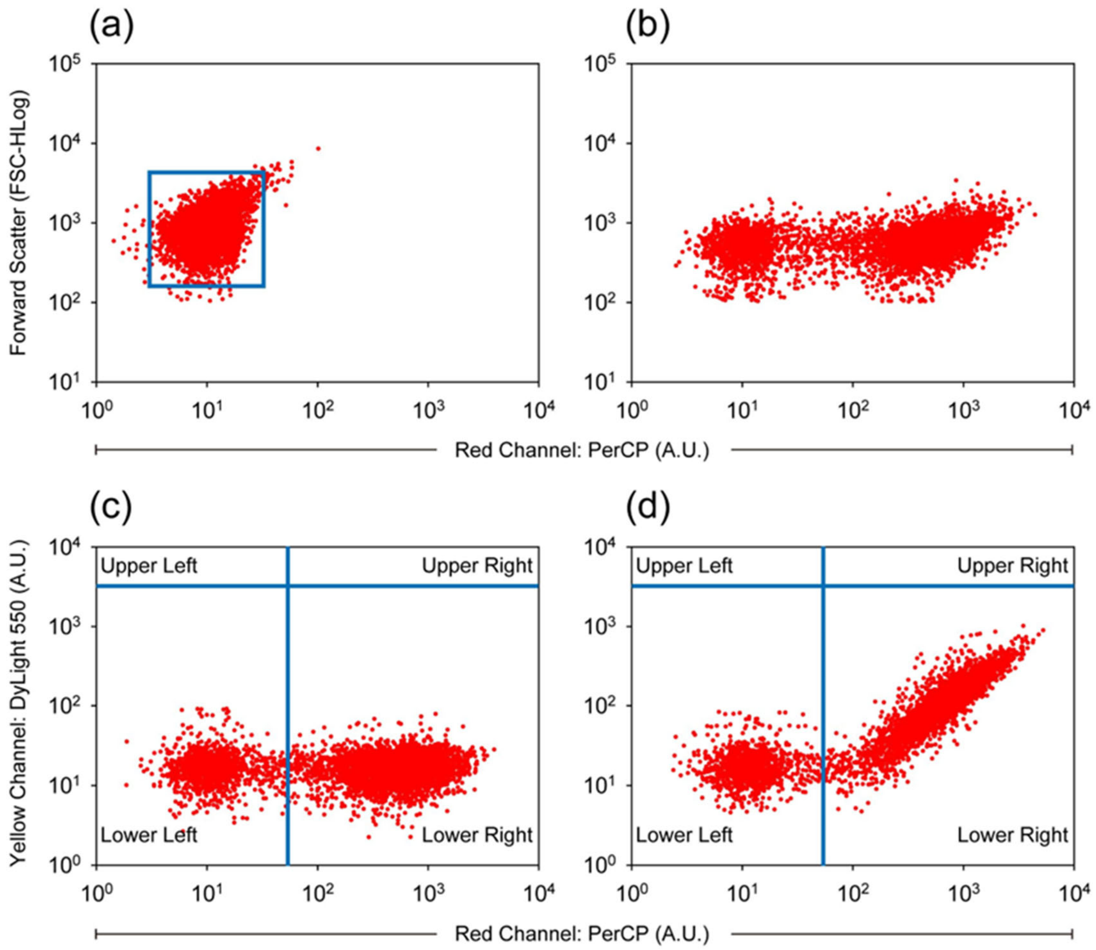


**Supplementary Fig. 1 | Example of the gating strategy for flow cytometry experiments. a**, Example gating strategy for analyzing yeast cells. The experiments were performed using a single type of non-stained yeast cells, which gave rise to a tight distribution on FSC vs red fluorescence plot. The gate in blue was used for subsequent analysis of fluorescent population. **b**, Example of FSC vs red fluorescence plot for Mb(S1)-expressing yeast cells stained with PerCP to assess the monobody expression. **c-d**, Example gating strategy for target binding experiments. For these experiments, Mb(S1)-expressing yeast cells were incubated in the absence (c) or the presence (d) of FtsZ, followed by staining with PerCP and DyLight 550 to assess the monobody expression and target binding, respectively. Binding of Mb(S1) to FtsZ was measured by taking the median fluorescence intensity (MFI) of 75–90% of the DyLight 550 fluorescent population in the lower right quadrant. For panel (d), the target protein was EcFtsZ at the concentration of 1,250 nM.

**Supplementary Video 1 | CryoEM map and model of KpFtsZ–Mb double helical tube.**
